# Inducible degradation-coupled phosphoproteomics identifies PP2A^Rts1^ as a novel eisosome regulator

**DOI:** 10.1101/2023.10.24.563668

**Authors:** Andrew G. DeMarco, Marcella G. Dibble, Mark C. Hall

## Abstract

Reversible protein phosphorylation is an abundant post-translational modification dynamically regulated by opposing kinases and phosphatases. Protein phosphorylation has been extensively studied in cell division, where waves of cyclin-dependent kinase activity, peaking in mitosis, drive the sequential stages of the cell cycle. Here we developed and employed a strategy to specifically probe kinase or phosphatase substrates at desired times or experimental conditions in the model organism *Saccharomyces cerevisiae.* We combined auxin-inducible degradation (AID) with mass spectrometry-based phosphoproteomics, which allowed us to arrest physiologically normal cultures in mitosis prior to rapid phosphatase degradation and phosphoproteome analysis. Our results revealed that protein phosphatase 2A coupled with its B56 regulatory subunit, Rts1 (PP2A^Rts1^), is involved in dephosphorylation of numerous proteins in mitosis, highlighting the need for phosphatases to selectively maintain certain proteins in a hypophosphorylated state in the face of high mitotic kinase activity. Unexpectedly, we observed elevated phosphorylation at many sites on several subunits of the fungal eisosome complex following rapid Rts1 degradation. Eisosomes are dynamic polymeric assemblies that create furrows in the plasma membrane important in regulating nutrient import, lipid metabolism, and stress responses, among other things. We found that PP2A^Rts1^-mediated dephosphorylation of eisosomes promotes their plasma membrane association and we provide evidence that this regulation impacts eisosome roles in metabolic homeostasis. The combination of rapid, inducible protein degradation with proteomic profiling offers several advantages over common protein disruption methods for characterizing substrates of regulatory enzymes involved in dynamic biological processes.

## INTRODUCTION

Reversible protein phosphorylation, catalyzed by the opposing activities of kinases and phosphatases, is a ubiquitous regulatory mechanism in eukaryotic organisms. Virtually all biological processes rely on some level of control by dynamic protein phosphoregulation. The cell division cycle is a well-studied example. At its core, cyclin-dependent kinase activity, driven by waves of cyclin expression and degradation, triggers key cell division events, including cell cycle entry, genome replication, and initiation of the mitotic state (1). In general, protein kinase activity rises from cell cycle start through metaphase of mitosis (1–4). Protein phosphatase activity, on the other hand, is generally suppressed in the early stages of mitosis and stimulated as cells enter anaphase (2, 4–7). Many proteins are maximally phosphorylated in mitosis, often nearly stoichiometrically (8). However, cells must maintain low phosphorylation states on some proteins during mitosis in the face of high kinase activity, which requires protein phosphatases acting in a highly specific manner on distinct substrates.

Here we describe a methodology to address this and other questions of dynamic protein phosphoregulation that couples rapid, inducible protein degradation with quantitative LC-MS/MS-based phosphoproteomics. We used the auxin-inducible degradation (AID) technology (9) in *S. cerevisiae* to achieve degradation of different components of the abundant protein phosphatase 2A (PP2A) within minutes in arrested mitotic cells and measured the corresponding changes in the phosphoproteome. This AID-based workflow has advantages over other genetic approaches for suppressing target protein function. Compared to methods like gene deletions, transcriptional repression, and RNA interference, AID is very rapid, on the same time scale achieved by specific chemical inhibitors. Moreover, using AID maintains natural promoter control of the target gene and normal cell physiology until the time of auxin addition. In principle, AID should minimize non-specific effects caused by permanent loss-of-function mutations or long-term repression of gene expression (10, 11) and should be superior at specifically revealing direct substrates of regulatory enzymes like kinases and phosphatases. Here, it allowed us to induce metaphase arrest in physiologically normal cells before target degradation, ensuring that any detected phosphoproteome changes could be attributed to the mitotic activity of the targeted phosphatase.

PP2A is a member of the phosphoprotein phosphatase superfamily, and comprises a group of heterotrimeric protein phosphatases sharing common catalytic (C) and scaffold (A) subunits with individual enzymes distinguished by unique regulatory (B) subunits that provide substrate specificity and regulation (6, 12, 13). PP2A enzymes are responsible for a large fraction of total phosphatase activity in eukaryotic cells and play diverse roles in numerous biological processes (6, 13, 14). Mammalian species have many B subunit genes and isoforms, falling into 4 families (7, 12). The two major classes, B55 and B56, have 4 and 5 isoforms, respectively, with unique functions in the cell cycle (7). *S. cerevisiae* has fewer B subunit genes (14), with a single B55 homolog (*CDC55*) and a single B56 homolog (*RTS1*), making *S. cerevisiae* convenient for individually studying the substrate specificity and biological function of these two major PP2A holoenzyme families. During mitosis in many eukaryotes, PP2A-B55 enzymes are potently inhibited by the MASTL/Greatwall kinase via the small endosulfine proteins to help promote the high protein phosphorylation required to establish the mitotic state (15–19). However, PP2A-B56 enzymes are unaffected by MASTL/Greatwall and maintain activity during mitosis (6, 7, 20–23). For example, in *S. cerevisiae,* PP2A^Rts1^ activity is essential for proper function of the spindle position checkpoint and to protect sister chromatid cohesion prior to anaphase (6, 23, 24). We therefore used AID-coupled phosphoproteomics to identify proteins whose phosphorylation state is actively regulated by PP2A^Rts1^ in mitosis. This method revealed that PP2A^Rts1^ regulates the phosphorylation and plasma membrane association of eisosomes, fungal-specific protein complexes that play important roles in nutrient import, lipid metabolism, and stress responses (25–28).

## EXPERIMENTAL PROCEDURES

### Strain and plasmid construction

Complete lists of strains and plasmids are provided in Tables S1 and S2, respectively. YAK201 was a generous gift from Dr. Ann Kirchmaier (Purdue University). YKA1202 was constructed by PCR-mediated integration of KanMX to replace the *ARG4* coding sequence. The lysine and arginine auxotrophy of YKA1202 and its derivative strains enables SILAC metabolic labeling (29) as another option for quantitative proteomics. Strains YKA1223, YKA1224, and YKA1225 were constructed by PCR-mediated integration of the 3xV5/IAA17 degron tag from pAR1099 at the 3’ end of the *CDC55*, *TPD3,* and *RTS1* genes, respectively, in YKA1202. These strains were further engineered by integrating *StuI-*linearized pAR1103, containing the *Oryza sativa* transport inhibitor response 1 (*OsTIR1*) gene expressed from the *ADH1* promoter, at the *leu2* locus to yield YKA1232-YKA1235. pAR1099 and pAR1103 were generous gifts from Dr. Adam Rudner (University of Ottawa). YKA1236 and YKA1238 were constructed by PCR-mediated integration of an EGFP tag with *TRP1* marker amplified from pYM29 (30) at the 3’ end of the *PIL1* gene into YKA1233 and W303, respectively. YKA1239 and YKA1240 were created by PCR-mediated integration of a *URA3* cassette amplified from pAG60 (31) to replace the *RTS1* coding sequence in YKA1238 and BY4741, respectively. All strains were verified by PCR and by immunoblotting when appropriate. pHLP735 and pHLP747 were constructed using the In-Fusion cloning system (Takara Bio). The *RTS1* gene, including 500 bp upstream of the start codon and the 3’ 3xV5 tag was amplified from YKA1233 genomic DNA and inserted into pRS413-GAL-ccdB-3xHA and pRS423-GPD-ccdB (32) that were cut with *EcoRV* and *Eco53KI* to remove the promoter-ccdB-3xHA sequence. The final plasmids were verified by WideSeq DNA sequencing.

### Yeast cell culture

Yeast cultures were grown in YPAD (10 g/L yeast extract, 20 g/L peptone, 40 mg/L adenine, 20 g/L dextrose) or synthetic dropout media (6.7 g/L yeast nitrogen base without amino acids, 20 g/L dextrose, 1.4 g/L nutrient dropout mix), where indicated, at 30°C with shaking at 225 rpm until the desired optical density at 600nm (OD_600_). For metaphase experiments, cultures were treated with 15 µg/ml nocodazole (Cayman Chemical) for ∼2.5 hours. Arrests were confirmed by microscopic inspection of cell morphology. For experiments using the AID system, cultures were split with one half being treated with 250 µM 3-indoleacetic acid (IAA; Millipore Sigma) and the other half with an equal volume of DMSO. For phosphoproteomic experiments trichloroacetic acid (TCA) was added to cultures at a final concentration of 10% and cells harvested by centrifugation. Cell pellets were washed twice with 10 mL 70% ethanol and twice with 10 mL 6 M urea. The final supernatant was removed, and pellets stored at -80°C until processing. Myriocin (Cayman Chemical), phytosphingosine (Enzo Life Sciences) and canavanine (Cayman Chemical), were included in liquid or solid growth media at the indicated concentrations. For agar plate growth assays, saturated overnight cultures were diluted to a starting OD_600_ of 1 then serially diluted in eight-fold steps in sterile PBS. 5 µl aliquots were spotted on agar plates, which were incubated up to 5 days at 30°C. For liquid growth assays, cells were diluted to a final OD_600_ of 0.01 in SD-arg media with and without canavanine. Growth was monitored for 60 hour at 30°C by measurement of OD_600_ every 15 minutes with constant orbital rotation in a BioTek Cytation1 plate reader. Data were analyzed using GraphPad Prism v10.0.

### Immunoblotting

Samples for immunoblotting were prepared as previously described (33). Samples were first separated on 10% SDS-PAGE gels in Tris-glycine-SDS buffer at 170V for 1 hour and transferred to 0.45 µm Immobilon-FL-PVDF (Millipore Sigma) or nitrocellulose (Bio-Rad) membranes in Tris-glycine buffer at constant 1.0 A for 1 hour using a Bio-Rad Trans-Blot Turbo system. Membranes were blocked with 5% non-fat dry milk in PBST. The following primary antibodies and dilutions were used overnight at 4 °C in PBST with 1% milk: mouse anti- V5 conjugated to horseradish peroxidase (HRP; 1:5,000; Invitrogen), mouse anti-V5 (1:5,000; Invitrogen), rabbit anti-G6PDH (1:5,000; Millipore Sigma), mouse anti-GFP (1:1,000; Roche). The following secondary antibodies and dilutions were used for 1 hour at room temperature: goat anti-rabbit StarBright Blue 700 (1:2,500; Bio-Rad), goat anti-rabbit IRDye 800 CW (1:5,000: LiCOR), goat anti-mouse-HRP and goat anti-rabbit-HRP (1:10,000; Jackson Laboratories). Blots using HRP-conjugated antibodies were developed with Clarity Western ECL Substrate (Bio-Rad). All blots were imaged on a ChemiDoc MP digital imager (Bio-Rad), and quantification of band intensities performed using ImageLab 6.1 (Bio-Rad). Relative abundances of AID target proteins were calculated by normalizing band intensities to the cognate loading control band and then to the untreated or time 0 sample intensity, where indicated.

### Fluorescence Microscopy

Pil1-EGFP was used as a marker for eisosome localization. For AID strains, IAA treatment lasted 1 hour. Cells were collected by centrifugation, washed with PBS, fixed in a 4% paraformaldehyde/4% sucrose solution for 10 minutes at room temperature, washed again with PBS, and stored at 4°C until imaging. Imaging was performed by mounting cells on poly-L-lysine coated slides. A series of Z-plane images was collected using the FITC channel at room temperature with a 100x oil objective on a confocal Nikon A1Rsi microscope. All images within an experiment were acquired with identical settings. Images were analyzed using ImageJ and statistical analysis was performed in Prism. Any adjustments were applied equally to all images within an experiment. Images shown are at the approximate mid-section of the cell.

### Phosphoproteomics sample preparation

Cell pellets were lysed with 0.5 mm zirconium oxide beads (Next Advance) in lysis buffer (7 M Urea, 50 mM Tris-HCl (pH 8.0), 50 mM sodium fluoride, 50 mM β-glycerophosphate, 1 mM MgCl_2_, 1 mM sodium orthovanadate, 10 µM pepstatin A, 10 µM bestatin) in a disrupter genie (Scientific Industries) until >90% lysis was achieved based on microscopic estimation. Extracts were clarified at 16,000 xg for 15 minutes. The soluble fraction was decanted and treated with 2.5 units/µl Benzonase® (Millipore Sigma) for 45 minutes at 30°C. Extracts were diluted to 1 M urea with 50 mM fresh ammonium bicarbonate. Tris(2-carboxyethyl)phosphine and chloroacetamide were added to final concentrations of 5 and 30 mM, respectively, and TrypZean® (Millipore Sigma) was added for a final 1:100 enzyme:protein ratio. The mixture was incubated for 24 h at 37°C, then quenched with 0.1% trifluoroacetic acid (TFA). On-column dimethyl stable isotope labeling was performed essentially as described (34) using Sep-Pak Vac 3cc (500 mg) C18 columns (Waters Corporation) with a vacuum manifold. Labeling reagent was 50 mM sodium phosphate buffer (pH 7.5), 0.6 M sodium cyanoborohydride (Alfa Aesar), and either 0.2% formaldehyde (CH_2_O; Neta Scientific) or deuterated formaldehyde (CD_2_O; Cambridge Isotope Laboratories). IAA-treated samples were labeled with CD_2_O and DMSO-treated samples with CH_2_O. Tryptic digests were loaded onto the C18 columns, labeling reagent applied, and columns washed 5 x 2 mL with 0.6% acetic acid and 5 x 2 mL with 0.1% TFA. Peptides were eluted with 60% acetonitrile in 0.1% TFA, the heavy and light labeled samples mixed in a 1:1 ratio based on bicinchoninic acid assay (Thermo Fisher), and the mixtures dried under vacuum.

### Phosphopeptide enrichment and fractionation

Strong cation exchange (SCX) phosphopeptide enrichment was adapted from (35, 36). Dried peptides were resuspended in 7 mM KH_2_PO_4_ (pH 2.7), 30% acetonitrile and 80 mM KCl and applied to an SCX spin column (Thermo Fisher) pre-equilibrated with 400 µL 7 mM KH_2_PO_4_ (pH 2.7) in 30% acetonitrile. After centrifugation at 2,000 xg for 5 minutes, the flow-through peptides were dried by vacuum centrifugation, resuspended in 0.1% TFA, desalted on 500 mg Sep-Pak Vac 3cc C18 columns as directed by the manufacturer and dried again.

Phosphopeptides were further enriched using PolyMAC (37) phosphopeptide kit (Tymora Analytical) as described by the manufacturer with the following adjustments. ∼10 mg of SCX pre-fractionated peptides were incubated with 350 µL of PolyMAC resin for 15 minutes at room temperature using end-over-end rotation. Phosphopeptides were eluted 3x with 50 µL PolyMAC elution buffer. Pooled elutions were dried under vacuum.

Phosphopeptides were then fractionated using a Pierce high pH reverse-phase peptide fractionation kit (Thermo Fisher) based on (38). Phosphopeptides were dissolved in 0.1% TFA for column loading and were sequentially step-eluted with 300 µL aliquots of 4%, 6%, 8%, 10%, and 20% acetonitrile in 0.1% triethylamine. Fractions were dried under vaccum.

### Immunoaffinity purification-mass spectrometry (IP-MS)

Immunoaffinity purification of Pil1- EGFP was performed as previously described (39, 40) with the following exceptions. *PIL1- EGFP* and untagged control cultures were grown to an OD_600_ of ∼0.8 and collected by centrifugation. Cells were lysed in 5 mL lysis buffer (150 mM potassium acetate, 20 mM HEPES (pH 7.4), 10% glycerol, 1% Triton X-100, 20 mM sodium fluoride, 50 mM β-glycerophosphate, 200 µM sodium orthovanadate) supplemented with 10 µM leupeptin, 1 mM benzamidine, 1 µM pepstatin, and 1 mM phenylmethylsulfonyl fluoride. Proteins were eluted with 80 µL of SDS dye without reducing agent by heating beads at 95 °C for 10 minutes and then subjected to SDS-PAGE. The entire lane was cut into pieces and subjected to in-gel digestion as described (41). Pooled extracted peptides were split. One portion was desalted on a 100 mg Sep-Pak column and analyzed directly by LC-MS/MS to identify interacting proteins. The second portion was desalted followed by on-column dimethyl labeling and phosphopeptide enrichment as described above. In this case, a third isotope state was generated using ^13^CD_2_O-formaldehyde and deuterated sodium cyanoborohydride (Cambridge Isotope Laboratories). The following labeling scheme was used: *PIL1-EGFP rts1*Δ “heavy”, *PIL1-EGFP RTS1* “medium”, and untagged control W303 “light”.

### Mass Spectrometry Data Acquisition

High pH C18 fractions were analyzed with a Dionex UltiMate 3000 RSLCnano System coupled to an Orbitrap Q Exactive HF Mass Spectrometer (Thermo Fisher). Peptides were resuspended in 2% acetonitrile in 0.1% formic acid and loaded at 5 μL/minute onto a PepMap C18 trap column (Thermo Scientific; 5 μm 100 Å particle size; 300 μm ID × 5 mm), then separated on an Aurora UHPLC C18-packed emitter column (Ionopticks; 1.6 μm 120 Å particle size; 75 μm ID x 25 cm) at 200 nL/minute using an increasing gradient of acetonitrile from 6% to 21% over 80 minutes, to 40% over 20 minutes, then to 80% over 5 minutes with column temperature of 40 °C. The mass spectrometer was operated in positive ion mode and used a top 10 data-dependent acquisition method. Precursor ion fragmentation was accomplished by higher energy collision dissociation at a normalized collision energy setting of 30%. The resolution of the orbitrap mass analyzer was set to 120,000 Hz and 7,500 Hz for MS1 and MS2, respectively, with a maximum injection time of 50 ms for MS1 and 20 ms for MS2. Dynamic exclusion was set to 60 s. Scan ranges were 375-1,500 m/z for MS1 and 300-1,250 m/z for MS2.

In the IP-MS experiment the following changes were made. Tryptic peptides were analyzed with a Vanquish Neo UHPLC system coupled to an Orbitrap Exploris 480 mass spectrometer (Thermo Scientific). Peptides were separated using an increasing acetonitrile gradient from 2% to 25% over 80 minutes, to 45% over 25 minutes, then to 95% over 10 minutes. The dependent scan number was increased to 20, resolution was 60,000 Hz for MS1 and 15,000 Hz for MS2, and MS1 scan range was 350-1650 m/z.

### Mass spectrometry data analysis

Raw MS data files were analyzed with MaxQuant (MQ) 2.0.1.0 (42, 43) using the Uniprot *S. cerevisiae* protein database and the following search settings. “Multiplicity” was set to “2” or “3” depending on the experiment. In the AID phosphoproteomics experiments “Light labels” were set to “DimethyLys0” and “DimethylNter0,” and “Heavy labels” were set to “DimethyLys4” and “DimethylNter4”. In the Pil1-EGFP IP-MS experiment, LFQ mode was used to identify interacting proteins. For phosphopeptide analysis from Pil1-EGFP IP-MS “Light labels” were set to “DimethyLys0” and “DimethylNter0”, “Medium labels” were set to “DimethyLys4” and “DimethylNter4,” and “Heavy labels” were set to “DimethylLys8” and “DimethylNter8”. In all experiments, digestion was set to Trypsin/P with a maximum of 4 missed cleavages. “Variable modifications” included phosphorylation on serine, threonine, and tyrosine, and “Fixed modifications” included carbamidomethylation on cysteines with a maximum of 5 modifications per peptide. Variable phosphorylation was not used in the MQ search for Pil1-EGFP interacting proteins. For all MQ searches, orbitrap “First search peptide tolerance” was 20 ppm; “Main search peptide tolerance” was 6 ppm; “Min. peptide length” was 5; “Max. peptide mass (Da)” was 5,000, and “Match between runs” was enabled. All other settings were default values. For the global phosphoproteomics experiments, the modification-specific peptides files were loaded into Perseus v1.6.15.0 (44), reverse hits were removed, and unique peptides identified in two of three trials were retained. Peptides were divided into phosphopeptide and non-phosphopeptide groups. For each trial, the H/L ratios of all peptides were normalized to the raw median H/L ratio of unmodified peptides (which should be exactly 1.0) to correct for any errors in mixing of peptide samples. The average median H/L ratio of unmodified peptides before normalization was 1.03 ± 0.07, indicating that mixing accuracy was high. H/L ratios were log_2_-transformed and data visualized with Prism and R. All mass spectrometry data are available via ProteomeXchange (45) with identifier PXD044337.

### Experimental Design and Statistical Rationale

All proteomic experiments were performed in biological triplicate. Statistical assessment of significance was applied to all experiments using widely-accepted software. All other biological experiments were performed independently at least three times. For quantitative measurements, appropriate statistical analyses have been conducted and reported. For fluorescence microscopy experiments 100 cells were quantified per sample and the analyses were conducted in a blinded fashion.

## RESULTS

### Rapid phosphatase degradation using AID generates detectable phosphoproteome changes

Many phosphoproteins are under active, dynamic control by opposing kinase and phosphatase activities. Inhibition of a kinase or phosphatase can lead to rapid changes in the abundance of phosphosites under its active control (46), and measuring such changes can be useful for identifying substrates and biological functions. We first tested if inducible degradation of a phosphatase via the AID system can elicit a rapid phosphoproteome change detectable with LC-MS/MS by targeting the PP2A B55 subunit, Cdc55, which has many substrates in diverse biological processes (6, 47, 48). While conducting this work, two other groups also successfully targeted Cdc55 with AID technology in *S. cerevisiae* to measure phosphoproteome changes (49, 50). We tagged the 3’ end of the chromosomal *CDC55* gene with the coding sequence for a 3xV5 epitope followed by the auxin-binding domain (ABD) of the *Arabidopsis thaliana* IAA17 protein in a strain constitutively expressing the *Oryza sativa* Tir1 F-box protein from the *ADH1* promoter (9). We first assessed the timing of Cdc55-ABD degradation and phosphoproteome changes following treatment of log-phase cultures with 500 µM of the natural auxin indole-3-acetic acid (IAA). Maximal Cdc55-ABD degradation occurred within 10 minutes of IAA treatment based on immunoblotting (**Fig. 1A**). In a preliminary experiment, we split a culture and treated half with IAA for 15 minutes and compared phosphopeptide abundances from the treated and untreated cells using dimethyl stable isotope labeling, phosphopeptide enrichment with PolyMAC, and data-dependent LC-MS/MS. However, we observed no increases in phosphopeptide abundances relative to unmodified peptides (**Fig. 1B**). We suspected there could be a lag in the appearance of phosphoproteome changes relative to target protein degradation observed by immunoblotting and, therefore, repeated this analysis with cultures harvested at 20, 40, and 60 minutes after IAA treatment. After 20 minutes, phosphopeptides exhibiting >2-fold increased abundance in the IAA-treated sample were identified, with a general shift in the phosphopeptide distribution towards increased abundance evident in the chromatogram (**Fig. 1C-D and Supplemental Dataset Sheet 1**), consistent with the expected broad impact of PP2A^Cdc55^ on protein phosphorylation. The fraction of upregulated phosphopeptides and shift in abundance distribution increased more after 40 minutes but did not change further in the 60-minute sample. We conclude that a modest delay exists between maximal Cdc55-ABD degradation and consequent changes to the phosphoproteome, but that the AID system can be used in conjunction with a phosphoproteomic workflow to reveal phosphosites under dynamic control of a target phosphatase or kinase.

**Figure 1.**
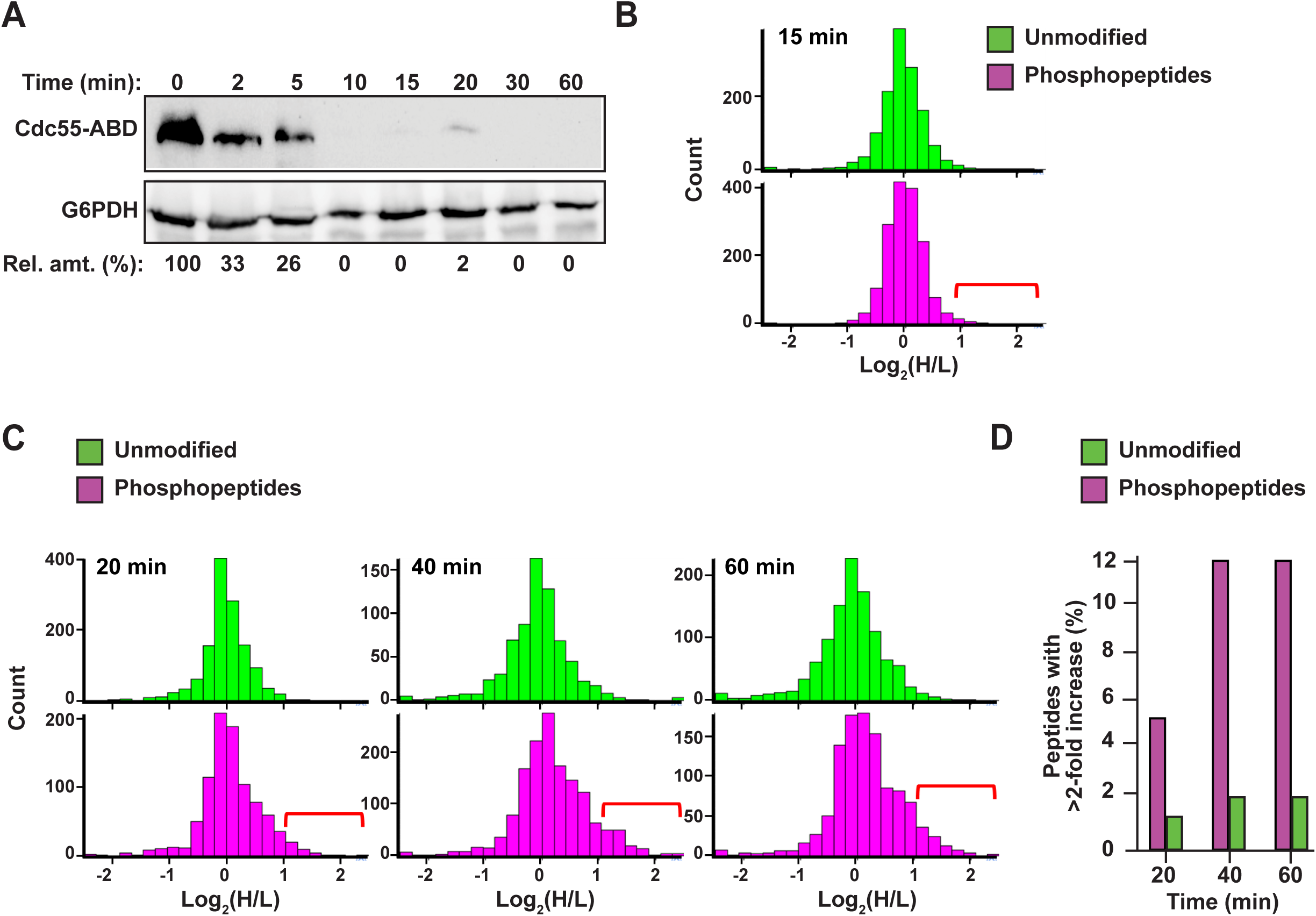
Auxin-induced phosphatase degradation causes detectable phosphoproteome changes. *A*, Cdc55-ABD level was monitored by immunoblotting with anti-V5 antibody at the indicated times after adding 500 µM IAA to log phase cultures. G6PDH is a loading control. “Rel. amt.” is the percentage of Cdc55-ABD remaining at each time relative to the starting 0 min sample. *B,* distribution of H/L ratios for identified unmodified peptides and phosphopeptides following 15-minute treatment of log phase *CDC55-ABD* culture with 500 µM IAA. Heavy (H)- labeled peptides are from the IAA-treated culture, and light (L)-labeled peptides are from mock-treated culture. Red bracket indicates the region where peptides with Cdc55-dependent phosphosites with increased H/L are expected to accumulate. *C,* same as panel B with cultures treated with IAA for 20, 40, or 60 minutes prior to quenching and harvesting. *D,* the fractions of identified phosphopeptides and unmodified peptides from panel C with >2-fold increase in IAA-treated *CDC55-ABD* culture were plotted as a function of time after IAA addition.

These preliminary experiments revealed the need to improve phosphopeptide enrichment and phosphoproteome coverage, so we further optimized our experimental workflow. We arrived at the workflow in **Fig. 2A** (see details in Methods). The new workflow had two notable improvements. First, we used an SCX bulk phosphopeptide enrichment immediately after the dimethyl labeling step, which removed a significant fraction of unphosphorylated peptides and improved the final ratio of phosphopeptides to unmodified peptides (**Fig. S1A-B)**. Second, we fractionated the final pool of PolyMAC-enriched phosphopeptides by high pH C_18_ chromatography to improve phosphoproteome coverage (**Fig. S1C**). Using this workflow we were routinely able to identify >10,000 peptides with typical phosphopeptide enrichment of 60- 80%. The remaining unmodified peptide fraction provided a useful representation of the general proteome for comparative purposes.

**Figure 2.**
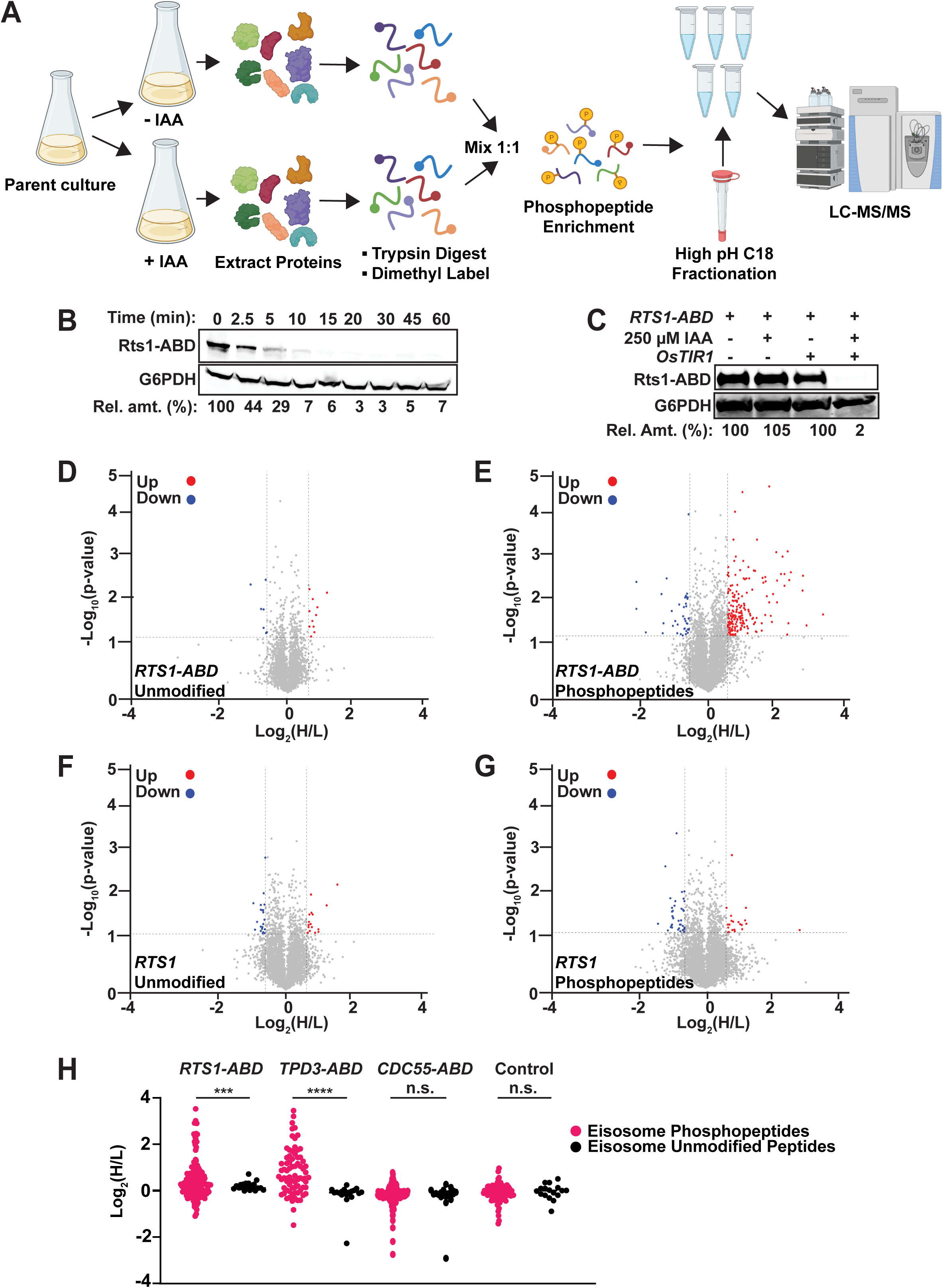
AID-mediated Rts1 degradation in mitotic cells causes rapid increase in phosphorylation of eisosome subunits and other proteins. *A,* Final AID-phosphoproteomics workflow. Parent culture expresses 1) target of interest tagged with ABD degron at its natural locus and 2) *OsTIR1* from the *ADH1* promoter. The phosphopeptide enrichment step includes SCX, followed by PolyMAC. IAA treatment is for 35 minutes. See Methods for full details. *B,* kinetics of Rts1-ABD degradation after treatment with 250 µM IAA in log phase cultures was monitored by immunoblotting with anti-V5 antibody. *C,* dependence of Rts1-ABD degradation on both *OsTIR1* and IAA was measured by anti-V5 immunoblotting. In B and C, G6PDH is a load control and “Rel. amt.” is either the percent Rts1-ABD remaining relative to time 0 (B) or relative to the -IAA sample (C). *D-G,* volcano plots of the Log_2_(H/L) ratios and associated p-values of unmodified peptides (D) and phosphopeptides (E) from the *RTS1-ABD* experiment and unmodified peptides (F) and phosphopeptides (G) from the untagged *RTS1* experiment. Regulated peptides were defined as -Log_10_(p-value) > 1.3 (p-value ≤ 0.05) and Log_2_(H/L) either ≤ -0.585 (down-regulated, blue) or ≥ 0.585 (up-regulated, red). Heavy dimethyl label (H) = IAA-treated; Light dimethyl label (L) = mock-treated. p-values were determined by t-test in Perseus. *H,* Log_2_(H/L) ratio distribution of all identified and quantified eisosome subunit phosphopeptides (pink) and unmodified peptides (black) from *RTS1-ABD, TPD3-ABD, CDC55-ABD,* and untagged control phosphoproteomic experiments in nocodazole-treated cultures. Statistical significance between average phosphopeptide and unmodified peptide Log_2_(H/L) values in each experiment was evaluated using a t-test, with p-value ranges defined as *** = p<0.001; **** = p<0.0001; n.s. (not significant) = p≥0.05.

Having established suitable conditions and workflow for measuring rapid phosphoproteome changes following phosphatase degradation, we turned to mitotic cells to address the question of phosphatases actively restraining protein phosphorylation by high mitotic kinase activities. We tagged PP2A’s B56 subunit, Rts1, with the same ABD degron for this purpose. Rts1 plays a key role in the response to osmotic stress and loss of Rts1 function renders *S. cerevisiae* hypersensitive to high salt (51, 52). *RTS1-ABD* cells were no more sensitive to 1 M NaCl than the parent strain in the absence of *OsTIR1,* but were unable to grow in the presence of *OsTIR1* and IAA, indicating that 1) the degron tag itself did not impair Rts1 function and 2) auxin-mediated degradation effectively reduced Rts1 function (**Fig. S1D**). 250 µM IAA was sufficient to reduce Rts1-ABD to ∼ 5% of its initial level within 15 minutes (**Fig. 2B, Fig. S1E**), dependent on the presence of OsTIR1 (**Fig. 2C**). We treated cultures of *RTS1-ABD* and its untagged parent strain with nocodazole to enrich for mitotic cells, split them, and treated half with 250 µM IAA, quenching with 10% TCA after 35 minutes (to account for the delay in phosphoproteome changes observed above). Rts1-ABD degradation and maintenance of mitotic arrest were confirmed (**Fig. S1F-G**), and phosphopeptide samples were prepared and analyzed from three biological replicates using our optimized workflow. We identified 10,781 phosphosites on 8,060 phosphopeptides from 1,596 proteins with PEP score (p-value) ≤0.05. We eliminated peptides identified in only a single trial for subsequent quantitative analyses (**Supplemental Dataset Sheet 2**). The mean ratio of heavy (+IAA, H) to light (mock-treated, L) dimethyl-labeled unmodified peptides across all three trials was 1.03 ± 0.07, indicating sample mixing was accurate. We set 1.5-fold change in H/L ratio as the cutoff for considering phosphopeptides regulated in a PP2A^Rts1^-dependent manner (Log_2_ values of -0.585 to +0.585) and required a p-value <0.05. With this threshold, only 0.36% of unmodified peptides qualified as upregulated upon degradation of Rts1 (**Fig. 2D**). In contrast, 3.3% of phosphopeptides (198 distinct peptides representing 233 total phosphosites on 111 proteins) were upregulated, while only 0.7% (42 peptides representing phosphosites on 33 proteins) were downregulated (**Fig. 2E**, **Supplemental Dataset Sheets 2-3**), consistent with the expected increase in substrate phosphorylation levels with reduced PP2A^Rts1^ activity. In the untagged control strain, only 0.4 % of phosphopeptides were upregulated and 0.6 % downregulated, similar to the fractions of unmodified peptides (**Fig. 2F-G, Supplemental Dataset Sheet 4**), indicating that most changes were dependent on loss of Rts1 and brief IAA treatment has little to no impact on the phosphoproteome.

The list of 111 proteins with upregulated phosphosites included several known PP2A^Rts1^ substrates, including Rck2 (51), Ace2 (53), Sgo1 (24), and Mad3 (22), and other candidate substrates identified in prior proteomic analyses of *rts1*Δ strains (**Table S3 and Fig. S1 H-I**). It also included many new candidate Rts1 substrates. These observations support the ability of the AID method to reveal PP2A^Rts1^ substrates and confirm that PP2A^Rts1^ actively counteracts phosphorylation of specific proteins during mitosis. We next looked for novel PP2A^Rts1^ substrate candidates in our data that might provide new insight into PP2A^Rts1^ biological functions during mitosis. We performed GO term and functional network analysis with STRING (54) using the 111 proteins with Rts1-dependent upregulated phosphosites (**Table S4 and Fig. S2**). The top-scoring GO category corresponded to a plasma membrane-associated protein complex termed the eisosome (Pil1, Seg1, and Eis1 proteins). The functional network analysis revealed a module with 6 proteins associated with the core eisosome complex (55) containing upregulated phosphosites (Pil1, Eis1, Msc3, Pkh2, Seg1, and Seg2). In fact, 20% of the upregulated phosphosites in our dataset came from these 6 eisosome subunits and 17 of the 50 most upregulated phosphopeptides are from Eis1, Seg1, and Seg2 (**Table 1, Supplemental Dataset Sheets 2-3**). Eisosomes are unique to fungi and interact with the plasma membrane to create long furrows that comprise a specialized region called the <u>M</u>embrane <u>C</u>ompartment of <u>C</u>an1 (MCC), which houses various nutrient transporters, lipid species, and membrane homeostasis regulators (25, 26, 28).

**Table 1.**
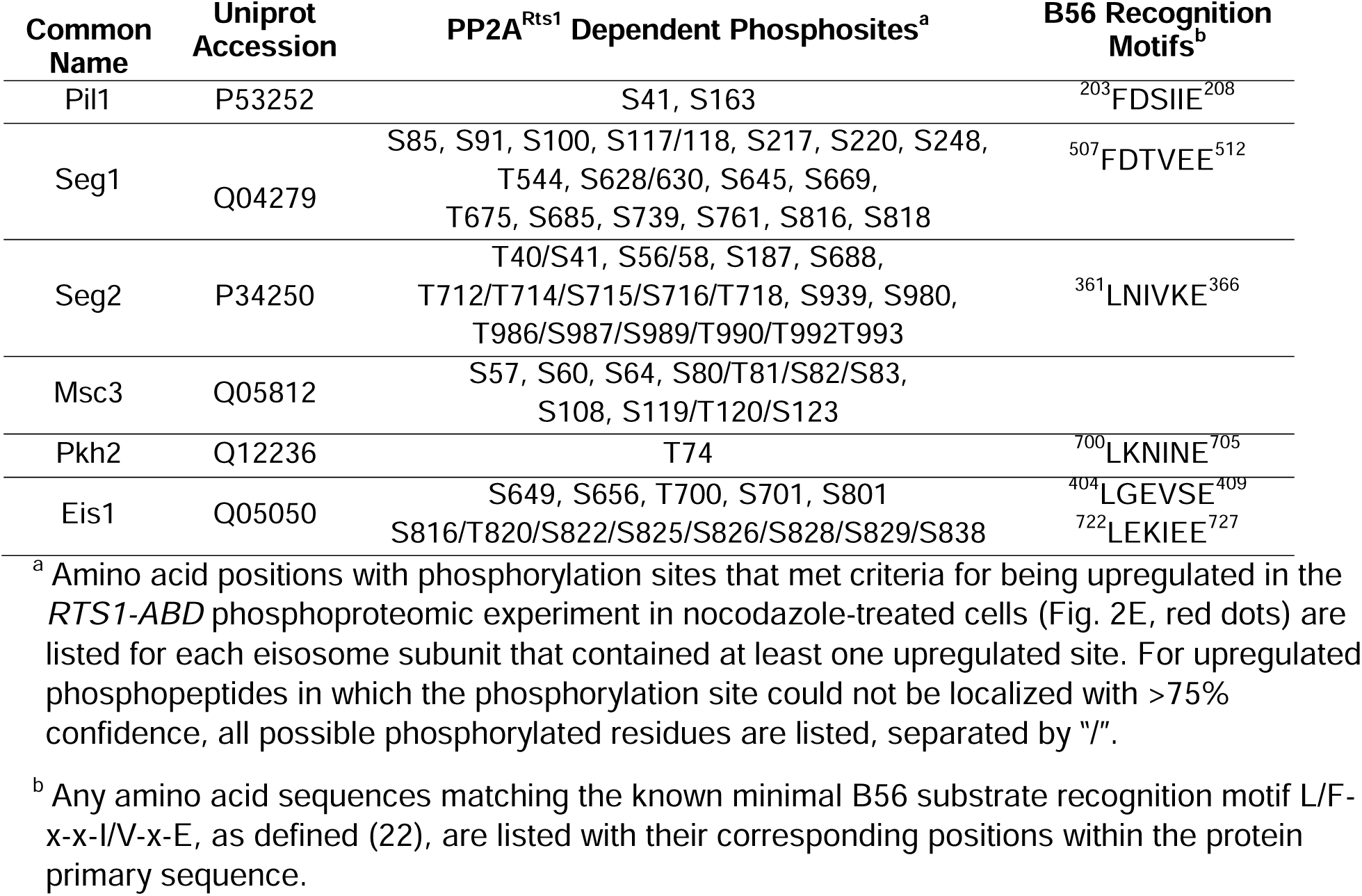
Rts1-regulated eisosome phosphosites.

To validate the effect of PP2A^Rts1^ on eisosome phosphorylation, we tagged the scaffolding subunit of PP2A, Tpd3, with the ABD degron and performed the same phosphoproteomic experiment under identical mitotic arrest conditions (**Fig. S3A-C)**. The *TPD3-ABD* data had greater variation in H/L ratio, so we used 2-fold change (log_2_ values of -1 to 1) as the threshold for considering peptides regulated in a Tpd3-dependent manner. Consistent with the PP2A^Rts1^ experiment, we observed significantly increased phosphorylation sites on the same 6 eisosome components after IAA-mediated Tpd3-ABD degradation (**Fig. S3D, Table S3, and Supplemental Dataset Sheet 5**). Both PP2A^Rts1^ and PP2A^Cdc55^ share the Tpd3 scaffold subunit. To determine if the effect on eisosome phosphorylation is specific to the Rts1 form of PP2A, we next performed an identical experiment with nocodazole-arrested *CDC55-ABD* cells. Variation in H/L ratio was more consistent with the *RTS1-ABD* experiment and we therefore used the same 1.5-fold change threshold for considering peptides regulated. Unlike our initial experiment in asynchronous log-phase cells, we detected very few significant phosphosite changes in metaphase-arrested cells following Cdc55 degradation, although there was a general shift towards increased H/L phosphopeptide ratios (**Fig. S3E-F**, **Supplemental Dataset Sheet 6**). This suggests that PP2A^Cdc55^ activity may be suppressed under the nocodazole arrest condition, consistent with the mitotic inhibition of PP2A-B55 enzymes in other systems (7). Importantly, no upregulated phosphopeptides were detected on eisosome subunits. The impacts of Rts1, Tpd3, and Cdc55 degradation on eisosome phosphosite abundance are summarized and compared in **Fig. 2H**, illustrating the specific effect of PP2A^Rts1^. B56 family proteins commonly recognize substrates by docking to the short linear motif [L/F]-X-X-[I/V]-X-[E] (22). The known PP2A^Rts1^ substrates Ace2, Rck2, and Mad3 all possess this motif, and mutations in the Rck2 and Mad3 motifs impaired Rts1 interaction and/or resulted in increased phosphorylation (22, 51). Five of the 6 identified eisosome-associated proteins in our dataset possess at least one occurrence of this B56 binding motif (**Table 1**), consistent with the possibility that eisosomes are directly dephosphorylated by PP2A^Rts1^. Overall, these data show that PP2A^Rts1^ specifically opposes eisosome subunit phosphorylation and might play a role in regulating eisosome function.

### PP2A^Rts1^ regulates Pil1 localization to the plasma membrane

The association of eisosomes with the plasma membrane is thought to be regulated by phosphorylation of eisosome subunits (26, 56). While kinases have been identified that influence eisosome- membrane interaction, including the PDK family members Pkh1 and Pkh2 that associate with eisosomes (56), protein phosphatases that regulate eisosomes have not been reported to our knowledge. Our results suggest that PP2A^Rts1^ could be involved in counteracting kinases that phosphorylate eisosomes and potentially in regulating eisosome association with the plasma membrane. The Pil1 core eisosome subunit is thought to bind membranes directly via its BAR domain (57, 58) and has been utilized frequently in fluorescence microscopy experiments as an indicator of eisosome assembly state at the plasma membrane (56–63). We therefore tagged *PIL1* with *EGFP* at its 3’ end in our *RTS1-ABD* strain to test if PP2A^Rts1^ influences eisosome association with the plasma membrane under the same conditions used in our phosphoproteomic experiments. Log phase cultures were incubated with nocodazole, then split and either treated with 250 µM IAA or mock-treated for 1 hour. Cells were fixed and the Pil1- EGFP fluorescence signal compared by confocal microscopy. We quantified both the number of eisosome puncta and the membrane:cytosol fluorescence ratio for 100 cells in each sample (**Fig. 3A, Fig. S4A**). Degradation of Rts1-ABD (**Fig. S4B)** resulted in a statistically significant (p<0.0001) decrease in membrane-associated fluorescence signal. The number of puncta per cell was only slightly reduced, but the difference was still statistically significant (p<0.01). These results suggest that PP2A^Rts1^ activity promotes eisosome assembly state at the plasma membrane in mitosis. We repeated the analysis in asynchronous log phase cultures and observed the same result, including a more pronounced impact on eisosome number (**Fig. 3B, Fig. S4C**). Thus, the effect of PP2A^Rts1^ on eisosome membrane association is not limited to mitosis. Finally, we evaluated the effect of *RTS1* gene deletion on the eisosome assembly state in asynchronous cultures. Consistent with the AID results, we observed large decreases in Pil1- EGFP membrane fluorescence and the number of puncta in the *rts1*Δ background, and this was complemented by reintroduction of *RTS1* on a plasmid (**Fig. 3C, Fig. S4D**). Taken together, these results indicate that PP2A^Rts1^ activity functions to promote Pil1 and eisosome association with the plasma membrane in dividing cells.

**Figure 3:**
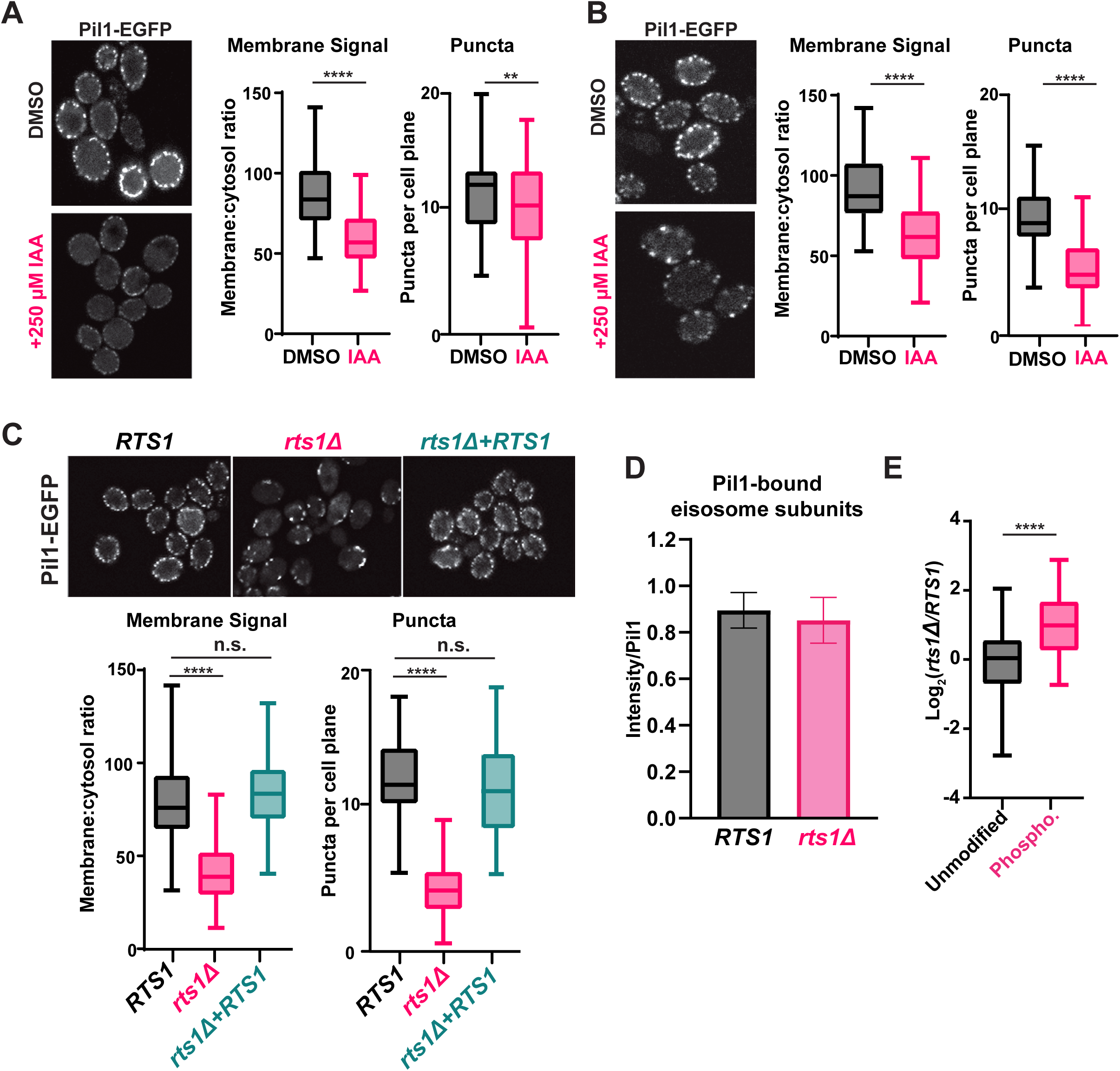
PP2A^Rts1^ promotes eisosome association with the plasma membrane but does not affect core eisosome subunit interactions. *A*, *RTS1-ABD PIL1-EGFP* cells were grown in YPAD and arrested in metaphase then split and treated with 250 µM IAA or an equivalent volume of DMSO for 1 hour. Fixed cells were imaged by confocal fluorescence microscopy. A single z-plane from the middle of the cells is shown. The membrane:cytosol fluorescence ratio (see Figure S4A) and number of puncta visible per cell were quantified from 100 cells of each sample, and the distributions plotted. The bars indicate the range of measured values, the boxes define the upper and lower quartiles, and the horizontal line inside the boxes is the median. T-tests were used to compare the IAA-treated and mock-treated samples, with p-values defined as follows: ** = p≤0.01; **** = p≤0.0001. *B,* same as A with asynchronous log-phase cultures. *C*, same as A with cells expressing *PIL1- EGFP* in the following backgrounds: *RTS1*, *rts1*Δ, or *rts1*Δ *+* CEN plasmid expressing *RTS1*. T-tests in C compared *RTS1* versus *rts1*Δ, and *RTS1* versus *rts1*Δ+*RTS1* plasmid. n.s. (not significant) = p≥0.05. *D*, IP-MS analysis of Pil1-associated eisosome subunits. The Log_2_- transformed intensities of peptides from all detected interacting eisosome subunits were divided by the summed Pil1-EGFP peptide intensity and used to create the bar and whisker plot. Results for the individual eisosome proteins are shown in Fig. S4F. *E*, the Pil1-normalized abundance ratios of unmodified peptides and enriched phosphopeptides (heavy label (H) = *rts1*Δ; light label (L) = *RTS1*) from the IP-MS experiment were plotted. T-tests were used to compare *RTS1* and *rts1*Δ in D, and the unmodified peptides vs. phosphopeptides in E with same p-value definitions described above.

Phosphorylation of the BAR domain proteins Pil1 and Lsp1 is generally believed to destabilize membrane-eisosome interaction (58), although other evidence suggests some phosphorylation sites may promote membrane assembly of eisosomes (59). The effect of Rts1 on Pil1-EGFP localization could therefore result from reduced membrane binding, but it could also have other explanations, including impaired oligomerization of Pil1/Lsp1, and/or reduced interaction with other eisosome components required for complex stability. To distinguish between these possibilities, we tested the effect of PP2A^Rts1^ on Pil1 protein interactions by IP-MS. Using conditions reported previously for IP-MS analysis of eisosomes (39, 40), we captured Pil1-EGFP from soluble protein extracts of *RTS1* and *rts1*Δ strains on anti-GFP antibody-coupled beads and performed LC-MS/MS analysis of the bound proteins. We detected five other core eisosome proteins in Pil1-EGFP IP samples but not in a control from the untagged parent strain (**Supplemental Dataset Sheet 7)**. We normalized each detected eisosome protein’s intensity to that of Pil1-EGFP and compared these ratios between *RTS1* and *rts1*Δ*. RTS1* status did not significantly affect Pil1 association with other eisosome components (**Fig 3D, Fig. S4E-F**). To confirm that *rts1*Δ had altered eisosome phosphorylation status in this experiment, we enriched phosphopeptides from a portion of the IP-MS samples, performed dimethyl labeling, and calculated the H/L (*rts1*Δ*/RTS1*) ratios for all the unique unmodified and phosphorylated peptides for the eisosome-associated proteins. Consistent with our AID phosphoproteome experiment, the average abundance of eisosome phosphopeptides was significantly higher in *rts1*Δ compared to *RTS1* (**Fig. 3E, Supplemental Dataset Sheet 7**). These data suggest that dephosphorylation of eisosomes by PP2A^Rts1^ primarily affects their interaction with the membrane but not the integrity of the complex. We also looked for Rts1- EGFP puncta at the membrane to determine if PP2A^Rts1^ might stably associate with membrane-bound eisosomes but saw no evidence for this (not shown).

### PP2A^Rts1^-mediated dephosphorylation of eisosomes plays a role in metabolic homeostasis

Our next goal was to determine if eisosome regulation by PP2A^Rts1^ is biologically important. We began by testing the role of PP2A^Rts1^ and eisosomes in sphingolipid metabolism because each has been separately implicated in regulation of a feedback loop involving TORC2 that controls sphingolipid homeostasis (40, 64–67). Both *rts1*Δ and *pil1*Δ strains were shown to be modestly resistant to myriocin, an inhibitor of serine palmitoyltransferase that catalyzes the first step in sphingolipid synthesis (66, 67). However, it is unknown if the Rts1 impact on this pathway is mediated through eisosomes. Using spot dilution assays we confirmed the slight resistance of *rts1*Δ to myriocin (**Fig. S5A**). However, this effect was not complemented by reintroduction of *RTS1* on a centromeric plasmid. In fact, expression of *RTS1* from a CEN plasmid led to stronger myriocin resistance. In contrast, the *RTS1* plasmid fully complemented the osmotic stress sensitivity of *rts1*Δ. This suggested that the myriocin resistance phenotype observed in our *rts1*Δ cells is not directly attributable to loss of *RTS1*. Furthermore, Rts1 degradation using the AID system had the opposite effect, strongly increasing sensitivity to myriocin (**Fig. 4A**). Given the inability to complement the *rts1*Δ phenotype by reintroducing *RTS1*, we focused only on our *RTS1-ABD* strain to test if PP2A^Rts1^ regulation of eisosomes is connected to sphingolipid metabolism.

**Figure 4:**
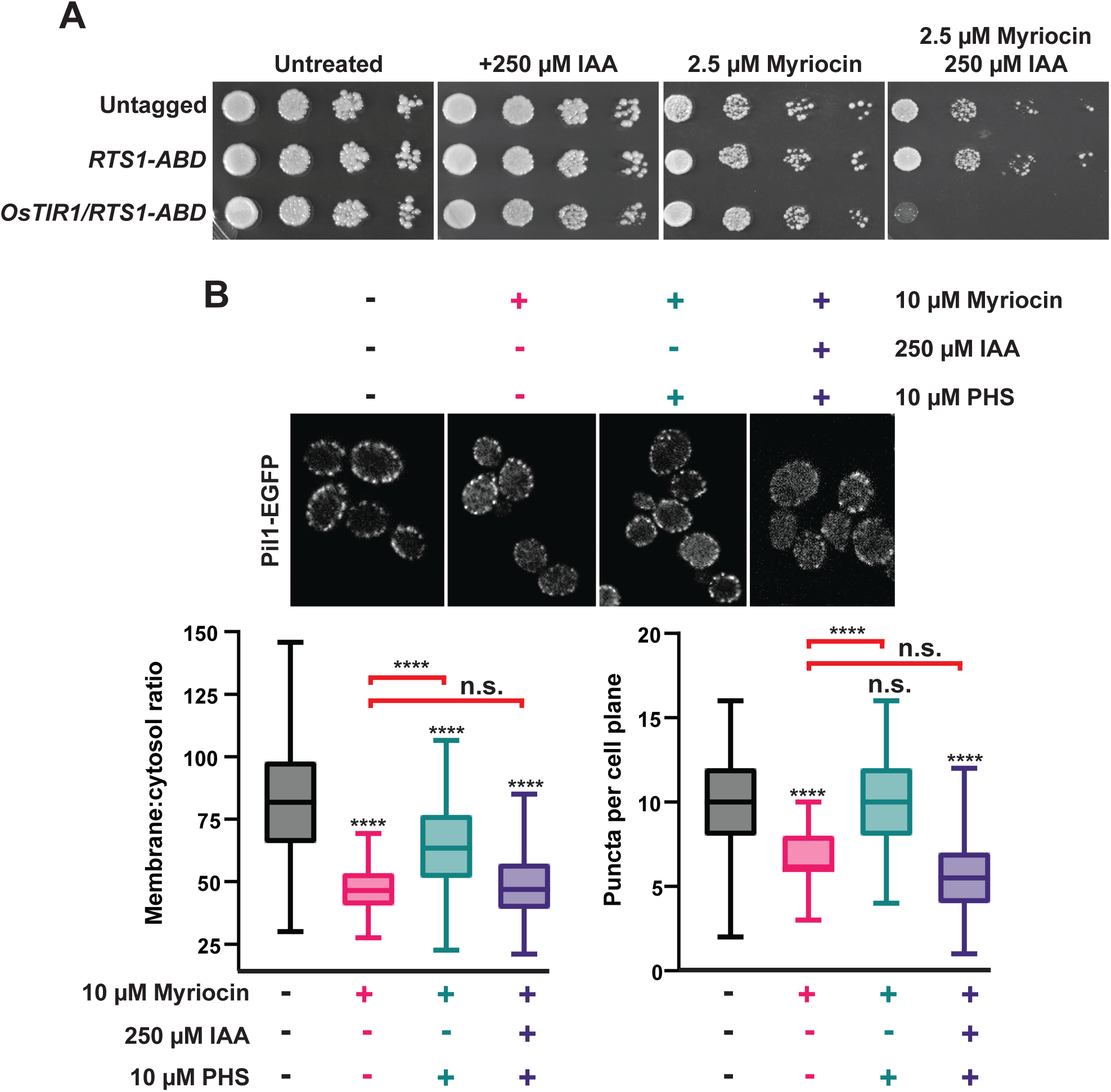
PP2A^Rts1^ influences eisosome-membrane interaction in response to sphingolipid levels. *A,* liquid cultures of the indicated strains were grown to saturation, serially diluted, and spotted on YPAD agar plates supplemented with myriocin and/or IAA, as indicated. Growth was monitored at 30°C for 48 hours for untreated and + 250 µM IAA plates and for 120 hours for plates containing myriocin. *B*, *RTS1-ABD PIL1-EGFP* cells were treated with the indicated combinations of 10µM myriocin (1 hour), 250 µM IAA or DMSO (35 minutes), and 10µM PHS (1 hour), in that order. Cells were fixed and imaged by confocal fluorescence microscopy. A single z-plane from the middle of the cells is shown. As before, the membrane:cytosol ratio and puncta per cell plane were quantified from 100 cells and differences statistically evaluated by t-tests, comparing each sample to the untreated control (**** = p<0.0001; n.s. (not significant) = p≥0.05). Red brackets indicate t-test results comparing the PHS-treated samples to myriocin alone.

To do this, we returned to confocal fluorescence microscopy with Pil1-EGFP. Eisosome association with the plasma membrane is reduced upon myriocin treatment and increased upon treatment with the sphingolipid precursor phytosphingosine (PHS), consistent with eisosomes playing a role in regulating sphingolipid synthesis in response to cellular need (40). To test if PP2A^Rts1^ affects eisosome-membrane association in response to changing sphingolipid levels, we treated a log phase *RTS1-ABD* culture with 10 µM myriocin for 1 hour to delocalize eisosomes from the membrane. Cultures were split and one half treated with 250 µM IAA to deplete Rts1. Both cultures were then treated with 10 µM PHS for 1 hour (**Fig. 4B, Fig. S5B**) to regenerate eisosomes. If our hypothesis were correct, recovery of eisosomes at the membrane in response to PHS should require dephosphorylation by PP2A^Rts1^. Consistent with prior work, myriocin treatment reduced the intensity and number of Pil1-EGFP puncta at the membrane and this was largely rescued by PHS treatment (**Fig. 4B**). Consistent with our hypothesis, degradation of Rts1 impaired recovery of Pil1-EGFP membrane fluorescence and puncta number in response to PHS treatment. This result suggests PP2A^Rts1^ regulation of eisosome localization may help cells respond to fluctuations of intracellular sphingolipid concentration.

We next explored if Rts1 might influence the nutrient transport function of eisosomes. Multiple amino acid and other nutrient transporters are specifically localized to the MCC defined by eisosomes (55). This includes the Can1 protein, a plasma membrane arginine permease, the loss of which confers resistance to the toxic arginine analog canavanine (68, 69). Loss of *PIL1* function also increases resistance to canavanine, presumably due to the reduced MCC area and number of Can1 symporters (68, 69). We hypothesized that PP2A^Rts1^ activity would also influence canavanine sensitivity by regulating the abundance of eisosomes at the membrane and, consequently, MCC size and number of active Can1 symporters. Indeed, in a serial dilution spot assay *rts1*Δ exhibited resistance to canavanine like deletions in eisosome subunit genes (**Fig. 5A**). Conversely, increasing *RTS1* copy number using CEN or 2µ plasmids increased sensitivity to canavanine (**Fig. 5B**). Importantly, increased canavanine sensitivity upon *RTS1* overexpression was not observed in a *pil1*Δ background. We further tested this hypothesis in liquid growth assays. Similar to the spot assays, increasing *RTS1* dose decreased the growth rate of cells with wild-type eisosome function in the presence of canavanine (**Fig. 5C**). In contrast, *RTS1* dose had no impact on the growth rate of *pil*Δ cells. These results are most easily explained by PP2A^Rts1^ influencing nutrient import by dephosphorylating eisosomes to promote their membrane association, with corresponding growth of the MCC and increase in concentration of MCC-associated transport proteins.

**Figure 5:**
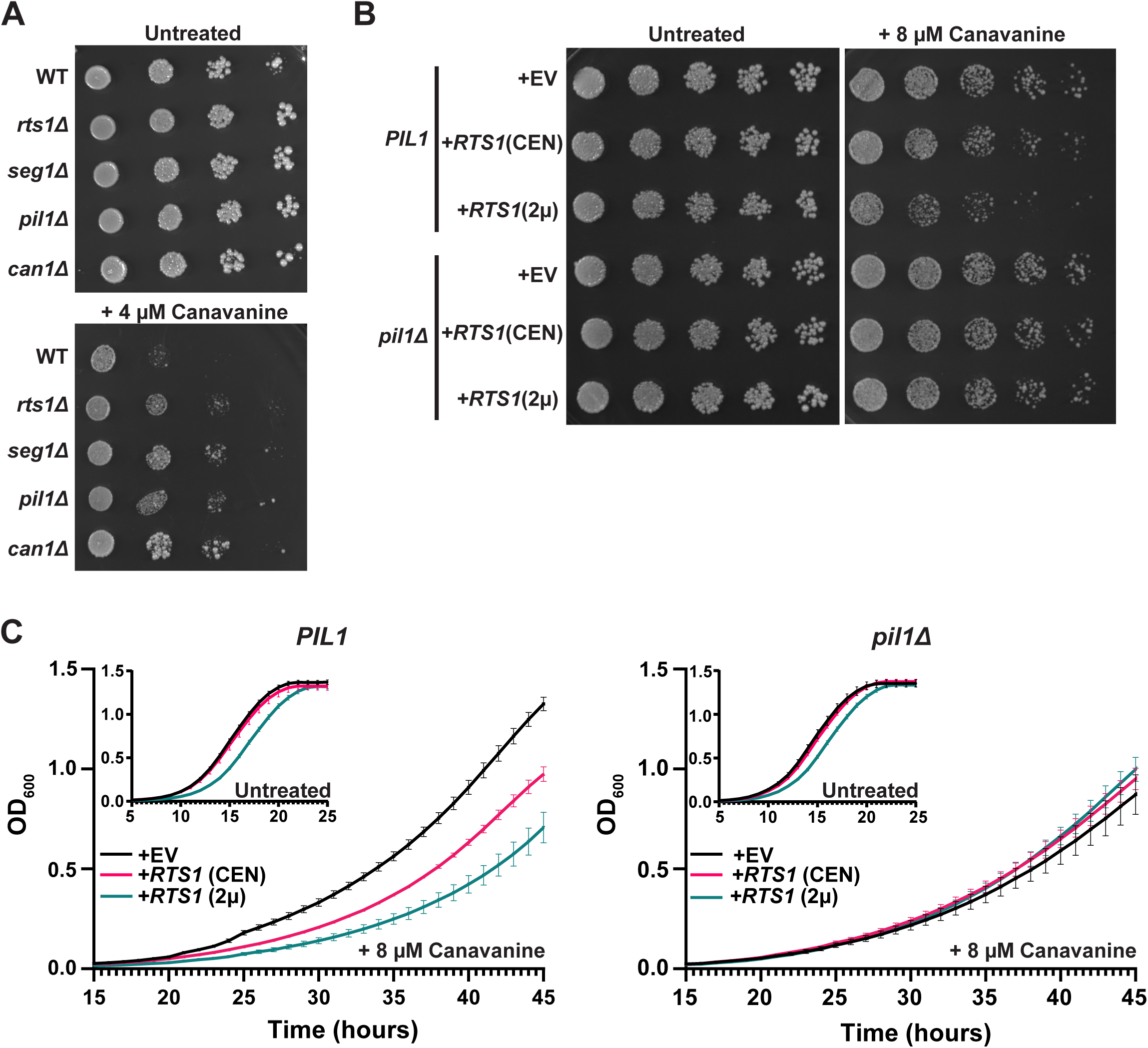
PP2A^Rts1^ regulation of eisosome localization is critical for the maintenance of amino acid homeostasis. *A,* Liquid cultures of the indicated strains were grown to saturation, serially diluted, and spotted on synthetic dropout agar plates lacking arginine and supplemented with canavanine. WT = wild-type. Growth was monitored at 30°C for 72 hours. *B,* same as A except wild-type *PIL1* and *pil1*Δ strains contained either empty vector (EV), or CEN or 2µ plasmids expressing *RTS1* from its natural promoter and growth was monitored at 30°C for 96 hours. In addition, starter cultures contained arginine to reduce canavanine toxicity and allow detection of increased canavanine sensitivity. Spot assays in A and B were performed 3 times with similar results. *C*, Liquid growth at 30 °C in a microplate with SD-arg media (with and without 8 µM canvanine) using strains from *B* was measured every 15 minutes as OD_600_. Data are the average of three independent cultures and error bars are standard deviations.

## DISCUSSION

Our primary goal was to develop a general workflow for combining the AID system with quantitative LC-MS/MS proteomics as a tool for characterizing substrates of regulatory enzymes like phosphatases and kinases. AID is a widely applicable inducible protein degradation technology, having been successfully employed in numerous model organisms in all major eukaryotic lineages outside plants. Its speed of action makes the system comparable to use of specific chemical inhibitors for characterizing global responses to loss of protein function, without the drawback of requiring a highly specific inhibitor for each protein of interest. The broad species applicability of AID, and its ease of use, make this proteomic workflow widely applicable. Moreover, AID offers advantages like maintaining natural target protein function until auxin addition, compared to the permanent, or longer-term, loss of function caused by gene deletions and RNA interference methods, respectively. This minimizes indirect effects on the proteome and should favor identification of direct substrates of the target (10, 11, 49). One drawback of this approach is the potential effect of the degron tag on target protein function, which is true for any fusion protein method. In addition, some targets may be refractory to robust degradation due to cellular localization effects and access of the degron tag to auxin. The AID- proteomics strategy is compatible with a variety of approaches for quantitation, including metabolic labeling and chemical labeling. It would be suitable for use with isobaric labeling reagents, like the tandem mass tag (TMT) system, to characterize time-resolved responses to stimuli or stress after target degradation. While we were conducting our work, other groups also demonstrated the ability to target phosphatases or kinases for degradation via the AID system and measure changes in the phosphoproteome in *S. cerevisiae* and human cells (49, 50, 70, 71).

We used our validated, optimized AID-phosphoproteomics workflow to identify proteins whose phosphorylation state is kept low in mitosis, when overall kinase activity is high, by the PP2A^Rts1^ phosphatase. We chose PP2A^Rts1^ for this purpose because it is known to be active during mitosis in *S. cerevisiae,* with functions including maintaining the spindle position checkpoint and centromeric cohesion prior to anaphase onset (23, 24). Our AID-mediated Rts1 degradation strategy identified known Rts1 substrates involved in mitotic processes (e.g. Ace2 (53), Rck2 (51), Mad3 (22), and Sgo1 (24)), as well as other candidate substrates. Our results also highlighted the ability of the AID-phosphoproteomics strategy to help distinguish the biological targets of different PP2A complexes. Surprisingly, a large fraction of Rts1-dependent phosphosites identified in our experiments were on core subunits of the eisosome complex. Eisosomes polymerize on the cytosolic face of the plasma membrane to create linear invaginations, or furrows, in the membrane. Eisosomes contain more than a dozen protein components (55), but Pil1, its paralog Lsp1, and Seg1 (and presumably its paralog Seg2) appear to comprise a critical structural core (39, 60, 72, 73). Pil1 and Lsp1 contain membrane-binding BAR domains and Seg1 promotes Pil1 association with the membrane, but detailed structural information on the complex and how it assembles into filaments along the membrane surface is still lacking. The furrows, termed “membrane compartment of Can1” (MCC) have a distinct phospholipid and protein composition, including selectively hosting a variety of nutrient transporters (55). MCCs also contain molecules that regulate lipid homeostasis, for example by controlling signaling through the TORC2 pathway (27). Eisosomes also appear to play important roles in response to certain stress conditions, have been linked to endocytosis (74), and are thought to be a reservoir of additional membrane material to allow for cell expansion (25). Eisosomes are regulated by phosphorylation of core subunits. The kinases Pkh1 and Pkh2 stably associate with the eisosome core and are at least partly responsible for its phosphorylation (56). While there is some debate about how phosphorylation affects eisosome structure and function, most evidence suggests that phosphorylation inhibits eisosome formation and it has been proposed that the negatively charged phosphate groups on the membrane interaction interface electrostatically impair membrane binding (57). Phosphatases that counteract eisosome phosphorylation have not been reported to our knowledge.

We provided evidence that PP2A^Rts1^ dephosphorylation of eisosomes is linked to known eisosome functions in regulating sphingolipid metabolism and amino acid transport. We first explored a potential connection to sphingolipid metabolism because both eisosomes and PP2A^Rts1^ have independently reported roles in control of sphingolipid synthesis via the TORC2 pathway (27, 64). In short, the MCC-associated proteins Slm1/2 are activators of TORC2 signaling and dissociation of eisosomes in response to low sphingolipid concentration was proposed to release Slm1/2 to bind and activate TORC2 to stimulate sphingolipid synthesis (75). Thus, eisosomes are negative regulators of TORC2 and sphingolipid synthesis in this model. Similarly, PP2A^Rts1^ was reported to be a negative regulator of TORC2-mediated sphingolipid synthesis, and consistent with this, *rts1*Δ strains were observed to be mildly resistant to the sphingolipid synthesis inhibitor myriocin (64, 66). Since PP2A activity is stimulated by ceramides (76, 77), intermediates in sphingolipid synthesis, this presented an attractive negative feedback model whereby ceramide accumulation would directly increase PP2A^Rts1^ activity to dephosphorylate eisosomes, increasing the MCC sequestration of Slm1/2 and consequent downregulation of TORC2. However, although we observed the same myriocin-sensitivity in our *rts1*Δ strains, this phenotype was not complemented by wild-type *RTS1*, suggesting it may be an indirect consequence of permanent Rts1 loss. Moreover, using our *RTS1-AID* strain, we found that IAA-induced Rts1-ABD degradation actually caused a strong sensitivity to myriocin. Thus, we were unable to generate support for this model. Nonetheless, the strong myriocin sensitivity after Rts1-ABD degradation confirms that PP2A^Rts1^ is important for responding to changes in sphingolipid levels. In support of this we then observed that the recovery of eisosomes following restoration of sphingolipid levels in myriocin-treated cells was dependent on Rts1. Thus, although the mechanism remains unclear, our results suggest PP2A^Rts1^ dephosphorylation of eisosomes plays a role in maintaining sphingolipid homeostasis.

We also provided evidence that PP2A^Rts1^ dephosphorylation of eisosomes is important for regulating import of nutrients like amino acids. Import of the arginine analog canavanine through the Can1 symporter, from which the MCC derives its name, is toxic. Cells lacking Can1 are resistant to canavanine, as are cells lacking eisosome subunits. This is explained by a loss MCC domains (and reduced Can1 concentration at membranes) when eisosomes are absent. We observed that *rts1*Δ cells were also resistant to canavanine, consistent with the idea that PP2A^Rts1^ increases eisosome and MCC abundance at the membrane. To directly test for a link between Rts1 and eisosomes in canavanine sensitivity, we increased Rts1 copy number and asked if the increased sensitivity to canavanine was dependent on eisosomes. It was, and although the phenotype itself was mild, the dependence on the Pil1 eisosome subunit was clear and reproducible. These results provided strong evidence that eisosome dephosphorylation by PP2A^Rts1^ is biologically important for controlling nutrient uptake through MCC-localized transporter proteins. Overall, we suggest that PP2A^Rts1^ is one of the primary phosphatases responsible for counteracting core eisosome phosphorylation to fine-tune the extent of eisosome and MCC assembly at the plasma membrane to maintain metabolic homeostasis and respond to changing metabolic needs.

Our Rts1-ABD phosphoproteomic dataset contains additional candidate mitotic substrates and potential biological functions for PP2A^Rts1^ that warrant future exploration. The STRING and GO term analyses revealed an enrichment of proteins related to cortical actin patches and endocytosis, including the network cluster centered around Ede1, Pan1, Vrp1, and Syp1 in Figure S2. Several of these proteins were also identified with Rts1-dependent phosphosites in proteomic studies of *rts1*Δ strains (48, 53). It is noteworthy that many of these proteins localize to the cell membrane like eisosomes, and eisosomes have been reported as cites of endocytosis (74). Another prominent cluster of functionally connected proteins, centered around Msn4, contains several transcription factors and protein kinases with functions in regulating nutrient metabolism and response to stress conditions. This cluster further supports the idea that PP2A^Rts1^ plays a broad role in regulating nutrient homeostasis.

In summary, we conclude that combining the AID system with LC-MS/MS-based proteomics is a powerful approach for characterizing the substrates and downstream effectors of regulatory enzymes involved in dynamic signaling pathways and other biological processes. We used it to demonstrate the importance of protein phosphatase activities like PP2A^Rts1^ to selectively restrain phosphorylation on key proteins during mitosis when overall cellular kinase activity is high. We expect that his general approach will be useful in characterizing additional phosphatases as well as many other regulatory enzyme classes in a wide range of organisms and biological contexts.

## Supporting information

Supplemental Proteomics Dataset

## ACKNOWLEDGEMENTS

MCH is supported by grants AI168050 and AI174123 from the National Institutes of Health, National Institute of Allergy and Infectious Diseases and this work was supported in part by an award from the Purdue University AgSEED program. The work was also supported by the Indiana Clinical and Translational Sciences Institute funded in part by Award Number UL1TR002529 from the National Institutes of Health, National Center for Advancing Translational Sciences, Clinical and Translational Sciences Award, and by pilot funding from the Purdue University Office of Research. The authors gratefully acknowledge the support for the Purdue Genomics Facility via the Purdue Institute for Cancer Research, NIH grant P30 CA023168. We thank Dr. Uma Aryal and Rodrigo Mohallem of the Purdue Proteomics Facility for assistance with proteomic data collection. We thank Dr. Xiaoguang Zhu and Shelly Tan of the Purdue Imaging Facility for assistance with confocal microscopy. We thank Dr. Elizabeth Tran for providing the GFP antibody.

## DATA AVAILABILITY

The mass spectrometry proteomics data have been deposited to the ProteomeXchange Consortium (http://proteomecentral.proteomexchange.org) via the PRIDE partner repository (45) with the dataset identifier PXD044337 and DOI 10.6019/PXD044337.

Other raw data, analyses and metadata will be made publicly available via the Purdue University Research Repository upon publication. Contact the corresponding author for access.

