## Supplemental Tables and Figures for "Inducible degradation-coupled phosphoproteomics identifies PP2A^Rts1^ as a novel eisosome regulator"

**Contents:**

Tables S1. Strains

Table S2. Plasmids

Table S3. Proteins with at least one Rts1-dependent upregulated phosphorylation site.

Table S4. STRING GO-Term analysis of proteins with at least one Rts1-dependent upregulated phosphorylation site.

Figures S1. Optimization and validation of AID phosphoproteomics method.

Figures S2. STRING protein functional network.

Figures S3. Mitotic AID phosphoproteomic analysis of *TPD3-ABD* and *CDC55-ABD*.

Figures S4. Supporting information for PP2ARts1 regulation of Pil1 localization and eisosome subunit interactions.

Figures S5. Supporting information for PP2A$^{Rts1}$ regulation of Pil1 localization in sphingolipid metabolism.

Supplemental Dataset. (separate Excel file with compiled proteomic results)

### Table S1. Strains

| Strain Name | Genotype | Source |
|---|---|---|
| W303 | *MATa ade2-1 his3-11,15 can1-100 leu2-3,112 trp1-1 ura3-1* | |
| BY4741 | *MATa his3Δ1 leu2Δ0 met15Δ0 ura3Δ0* | |
| *pil1Δ* | *BY4741 pil1::KanMX* | Horizon Discovery |
| *seg1Δ* | *BY4741 seg1::KanMX* | Horizon Discovery |
| *can1Δ* | *BY4741 can1:KanMX* | Horizon Discovery |
| YAK201 | *MATa lys2Δ his3-11,15 can1-100 leu2-3,112 trp1-1 ura3-1* | Ann Kirchmaier |
| YKA1202 | *MATa arg4::KanMX lys2Δ his3-11,15 can1-100 leu2-3,112 trp1-1 ura3-1* | This study |
| YKA1223 | *MATa CDC55-3xV5/IAA17:NatMX arg4::KanMX lys2Δ his3-11,15 can1-100 leu2-3,112 trp1-1 ura3-1* | This study |
| YKA1224 | *MATa TPD3-3xV5/IAA17:NatMX arg4::KanMX lys2Δ his3-11,15 can1-100 leu2-3,112 trp1-1 ura3-1* | This study |
| YKA1225 | *MATa RTS1-3xV5/IAA17:NatMX arg4::KanMX lys2Δ his3-11,15 can1-100 leu2-3,112 trp1-1 ura3-1* | This study |
| YKA1232 | *MATa TPD3-3xV5/IAA17:NatMX arg4::KanMX lys2Δ his3-11,15 can1-100 leu2::ADH1p-OsTIR1:LEU2 trp1-1, ura3-1* | This study |
| YKA1233 | *MATa RTS1-3xV5/IAA17:NatMX arg4::KanMX lys2Δ his3-11,15 can1-100 leu2::ADH1p-OsTIR1:LEU2, trp1-1, ura3-1* | This study |
| YKA1234 | *MATa CDC55-3xV5/IAA17:NatMX arg4::KanMX lys2Δ his3-11,can1-100,15 leu2::ADH1p-OsTIR1:LEU2, trp1-1, ura3-1* | This study |
| YKA1235 | *MATa arg4::KanMX lys2Δ his3-11,15 can1-100 leu2::ADH1p-OsTIR1:LEU2, ura3-1, trp1-1* | This study |
| YKA1236 | *MATa RTS1-3xV5/IAA17:NatMX arg4::KanMX lys2Δ his3-11,15 can1-100 leu2::ADH1p-OsTIR1:LEU2, ura3-1, trp1::PIL1-EGFP:TRP1* | This study |
| YKA1238 | *MATa ade2-1 his3-11,15 can1-100 leu2-3,112 trp1::PIL1-EGFP:TRP1 ura3-1,* | This study |
| YKA1239 | *MATa ade2-1 can1-100 his3-11,15 leu2-3,112 trp1::PIL1-EGFP:TRP1 ura3-1, rts1::URA3, can1-100* | This study |
| YKA1240 | *MATa his3Δ1 leu2Δ0 met15Δ0 ura3Δ0 rts1::URA3* | This study |

### Table S2. Plasmids

| Name | Yeast Origin | Promoter | Bacterial Marker | Yeast Marker | Expressed Protein | Source |
|---|---|---|---|---|---|---|
| pAR1103 | Integrating | *ADH1* | $Amp^R$ | *LEU2* | *Oryza sativa* Tir1 | Adam Rudner |
| pAR1099 | N/A | N/A | $Amp^R$ | *NatMX* | - | Adam Rudner |
| pRS413-GAL-ccdB-3xHA | CEN | N/A | $Amp^R$ | *HIS3* | N/A | (32) |
| pRS423-GPD-ccdB | 2µ | N/A | $Amp^R$ | *HIS3* | N/A | (32) |
| pHLP735 | CEN | *RTS1* | $Amp^R$ | *HIS3* | Rts1-3xV5 | This study |
| pHLP747 | 2µ | *RTS1* | $Amp^R$ | *HIS3* | Rts1-3xV5 | This study |

$Amp^R$ – β-lactamase gene, providing ampicillin resistance

NatMX – gene encoding resistance to nourseothricin

N/A – not applicable

**Table S3:** Proteins with at least one Rts1-dependent upregulated phosphorylation site

| Common name | Uniprot Accession | Overlap with Touati et al. (78) | Overlap with Zapata et al. (53) | Overlap with *TPD3-ABD* |
|---|---|---|---|---|
| ACE2 | P21192 | | Y | Y |
| ACM1 | Q08981 | Y | | |
| AFR1 | P33304 | | | |
| AIM21 | P40563 | | Y | Y |
| AIM3 | P38266 | | | Y |
| AIP5 | P43597 | | | Y |
| ASK10 | P48361 | | Y | |
| ASM4 | Q05166 | | | |
| AVO1 | Q08236 | | | |
| BCK2 | P33306 | | Y | |
| BNI4 | P53858 | | Y | Y |
| BNR1 | P40450 | | | |
| BOI2 | P39969 | | | |
| BUG1 | Q12191 | | | |
| BUL2 | Q03758 | | Y | |
| CIP1 | Q02606 | | | |
| CKI1 | P20485 | | | Y |
| CRN1 | Q06440 | | | |
| CRP1 | P38845 | | | |
| CYC7 | P00045 | | | |
| CYK3 | Q07533 | | | |
| DCS2 | Q12123 | | | Y |
| DEF1 | P35732 | | | |
| DRE2 | P36152 | | Y | Y |
| DSF2 | P38213 | Y | Y | |
| EAP1 | P36041 | | | Y |
| EDE1 | P34216 | Y | Y | Y |
| EIS1 | Q05050 | Y | | Y |
| ENT4 | Q07872 | Y | Y | Y |
| ENT5 | Q03769 | | | |
| EPO1 | P39523 | | | Y |
| ESC1 | Q03661 | Y | | |
| FUN19 | P28003 | Y | | Y |
| GAT1 | P43574 | | | |
| GCS1 | P35197 | | | Y |
| GIS1 | Q03833 | | | |
| GLN3 | P18494 | | | Y |
| HAA1 | Q12753 | | | |
| HRK1 | Q08732 | | | |
| HRP1 | Q99383 | | Y | |
| HSP42 | Q12329 | | | Y |
| ICS2 | P38284 | | | |
| ISF1 | P32488 | | | |
| JIP4 | Q03361 | | | |
| LRE1 | P25579 | Y | Y | |
| MAD3 | P47074 | | | |
| MBR1 | P23493 | | | Y |
| MFB1 | Q04922 | | | |
| MIF2 | P35201 | | | |
| MIX17 | Q03667 | | | Y |
| MLF3 | P32047 | Y | Y | Y |

| | | | | |
|---|---|---|---|---|
| MOT2 | P34909 | | | Y |
| MSC3 | Q05812 | | | Y |
| MSG5 | P38590 | | | |
| MSN4 | P33749 | | | Y |
| MSO1 | P53604 | | | |
| MUK1 | Q02866 | | | |
| NBA1 | Q08229 | | Y | Y |
| NRG1 | Q03125 | | | |
| NUP2 | P32499 | | | Y |
| OPY2 | Q06810 | | | |
| ORC6 | P38826 | Y | Y | |
| PAL1 | Q05518 | Y | | |
| PAL2 | P38809 | | | Y |
| PAN1 | P32521 | | Y | Y |
| PAR32 | Q12515 | | | Y |
| PBP1 | P53297 | | | Y |
| PDS1 | P40316 | | | |
| PET10 | P36139 | | | |
| PIB2 | P53191 | | | |
| PIL1 | P53252 | | | Y |
| PKH2 | Q12236 | | | Y |
| PRM5 | P40476 | | | |
| PSD2 | P53037 | | | |
| RCK2 | P38623 | | | Y |
| RCN2 | Q12044 | | | |
| RFM1 | Q12192 | | | |
| RIM15 | P43565 | | | |
| RTG1 | P32607 | | | |
| RTS3 | P53289 | | | Y |
| SAC7 | P17121 | | Y | |
| SEG1 | Q04279 | Y | | Y |
| SEG2 | P34250 | | | Y |
| SGM1 | P47166 | | | |
| SGO1 | Q08490 | | | |
| SIP1 | P32578 | | | |
| SIP5 | P40210 | | | |
| SIS2 | P36024 | | | |
| SKO1 | Q02100 | | Y | |
| SMY2 | P32909 | | | |
| SRO9 | P25567 | Y | | |
| SSZ1 | P38788 | | | |
| STP4 | Q07351 | | | Y |
| SWI5 | P08153 | | Y | Y |
| SYH1 | Q02875 | | Y | Y |
| SYP1 | P25623 | Y | | |
| TCO89 | Q08921 | | | |
| TDA11 | P38854 | | | |
| TOS7 | Q08157 | | | |
| TSL1 | P38427 | | | Y |
| UBX7 | P38349 | | | Y |
| UIP4 | Q08926 | Y | Y | Y |
| USV1 | Q12132 | | | |
| VAN1 | P23642 | | | |
| VHS2 | P40463 | | | Y |
| VRP1 | P37370 | | | Y |
| YER158C | P40095 | | | Y |

| | | | |
|---|---|---|---|
| YGR130C | P43597 | | |
| YLR257W | Q06146 | | |
| YMR295C | Q03559 | | Y |
| ZRG8 | P40021 | Y | |

**Table S4:** STRING GO-Term analysis of proteins with at least one Rts1-dependent upregulated phosphorylation site.

| #term ID | term description | Strength[a] | false discovery rate | Gene members from Rts1-ABD dataset |
|---|---|---|---|---|
| GO:0032126 | Eisosome | 1.34 | 0.031 | YGR086C,YMR031C,YMR086W |
| GO:0044615 | Nuclear pore nuclear basket | 1.34 | 0.031 | YAR002W,YDL088C,YLR335W |
| GO:0005844 | Polysome | 0.97 | 0.0496 | YCL037C,YGR178C,YHR064C,YKL204W |
| GO:0061645 | Endocytic patch | 0.89 | 0.0057 | YBL047C,YBR108W,YCR030C,YIR003W,YIR006C, YLL038C,YLR337C |
| GO:0032153 | Cell division site | 0.86 | 0.0375 | YCR030C,YDL117W,YDR348C,YIL159W,YNL233W |
| GO:0030863 | Cortical cytoskeleton | 0.85 | 0.0036 | YBL047C,YBR108W,YDL117W,YGR086C,YIR003W, YIR006C,YLL038C,YLR337C |
| GO:0030479 | Actin cortical patch | 0.83 | 0.0215 | YBL047C,YBR108W,YIR003W,YIR006C,YLL038C, YLR337C |
| GO:0030864 | Cortical actin cytoskeleton | 0.81 | 0.0107 | YBL047C,YBR108W,YDL117W,YIR003W,YIR006C, YLL038C,YLR337C |
| GO:0005935 | Cellular bud neck | 0.7 | 0.00027 | YAL041W,YBL047C,YCL051W,YCR030C,YDL117W, YDR348C,YER033C,YER114C,YIL117C,YIL159W, YIR006C,YLR337C,YNL233W,YNR049C,YOL070C |
| GO:0005938 | Cell cortex | 0.7 | 0.00067 | YBL047C,YBR108W,YCR030C,YDL117W,YDR348C, YDR389W,YGR086C,YIR003W,YIR006C,YLL038C, YLR337C,YNL233W,YOL100W |
| GO:0005937 | Mating projection | 0.7 | 0.0053 | YAL041W,YBL016W,YBL047C,YCR030C,YDR085C, YDR348C,YER033C,YIR003W,YIR006C,YLR337C |
| GO:0043332 | Mating projection tip | 0.7 | 0.0092 | YAL041W,YBL016W,YBL047C,YCR030C,YDR348C, YER033C,YIR003W,YIR006C,YLR337C |
| GO:0005934 | Cellular bud tip | 0.66 | 0.0428 | YAL041W,YBL047C,YBR007C,YCR030C,YDR348C, YER033C,YNR049C |
| GO:0030427 | Site of polarized growth | 0.63 | 0.00027 | YAL041W,YBL016W,YBL047C,YBR007C,YCL051W, YCR030C,YDL117W,YDR348C,YER033C,YER114C, YIL117C,YIL159W,YIR003W,YIR006C,YLR337C, YNL233W,YNR049C,YOL070C |
| GO:0005933 | Cellular bud | 0.62 | 0.00066 | YAL041W,YBL047C,YBR007C,YCL051W,YCR030C, YDL117W,YDR348C,YER033C,YER114C,YIL117C, YIL159W,YIR006C,YLR337C,YNL233W,YNR049C, YOL070C |
| GO:0005856 | Cytoskeleton | 0.6 | 0.00098 | YBL047C,YBR108W,YCR030C,YDL117W,YDL226C, YDR171W,YDR389W,YER114C,YGR086C, YIR003W,YIR006C,YLL038C,YLR337C,YNL233W, YOR073W |

[a] $Log_{10}$(observed/expected). Observed is the number of proteins in the data set which are annotated with the indicated GO-term. Expected is the number expected to contain this annotation in a random set of proteins from a list of identical length.

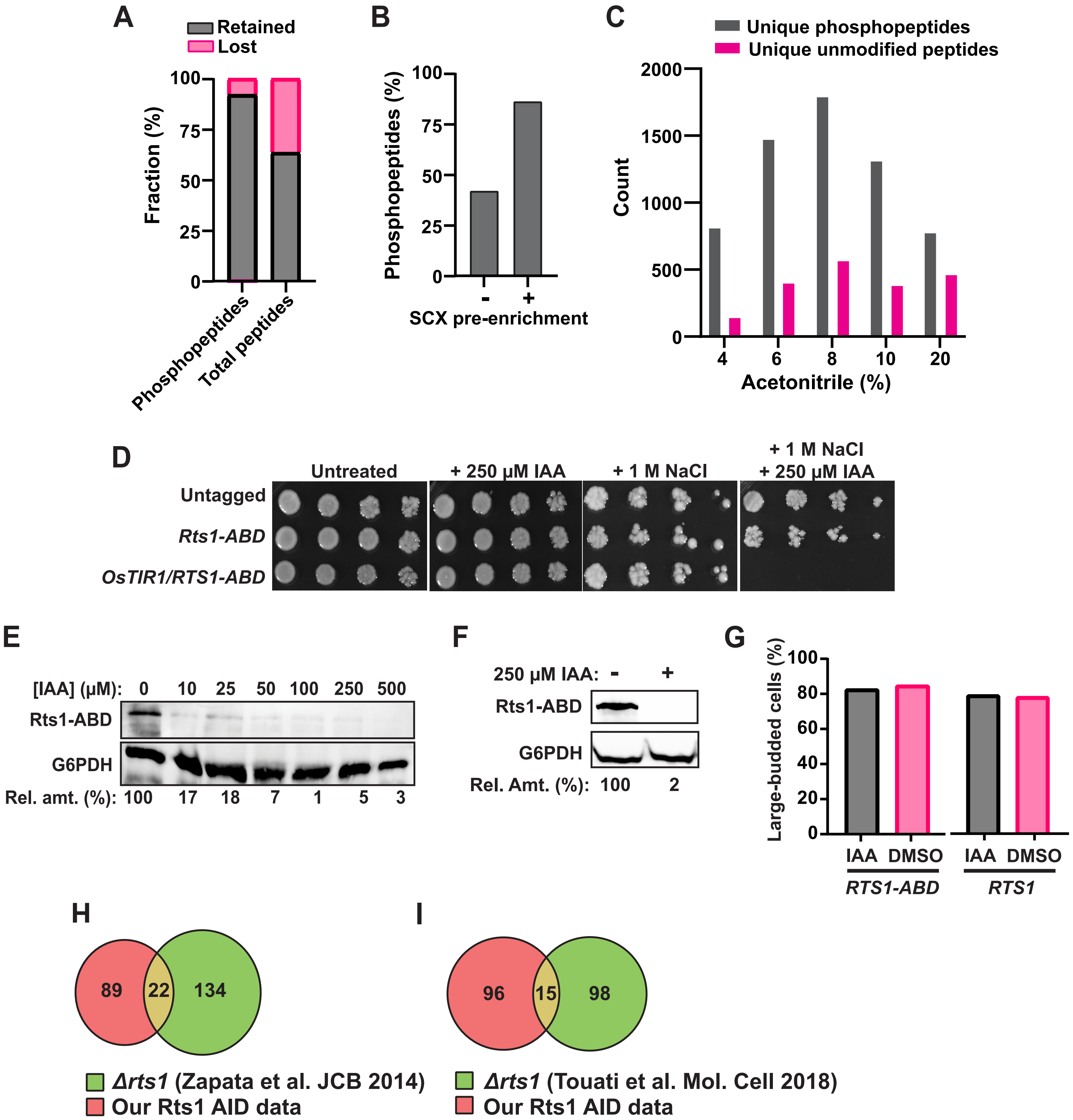
A
Retained
Lost
Fraction (%)
100
75
50
25
0
Phosphopeptides
Total peptides
B
Phosphopeptides (%)
100
75
50
25
0
-
+
SCX pre-enrichment
C
Unique phosphopeptides
Unique unmodified peptides
Count
2000
1500
1000
500
0
4
6
8
10
20
Acetonitrile (%)
D
Untreated
+ 250 µM IAA
+ 1 M NaCl
+ 1 M NaCl
+ 250 µM IAA
Untagged
Rts1-ABD
OsTIR1/RTS1-ABD
E
[IAA] (µM): 0 10 25 50 100 250 500
Rts1-ABD
G6PDH
Rel. amt. (%): 100 17 18 7 1 5 3
F
250 µM IAA: - +
Rts1-ABD
G6PDH
Rel. Amt. (%): 100 2
G
Large-budded cells (%)
100
80
60
40
20
0
IAA DMSO
RTS1-ABD
IAA DMSO
RTS1
H
89
22
134
Δrts1 (Zapata et al. JCB 2014)
Our Rts1 AID data
I
96
15
98
Δrts1 (Touati et al. Mol. Cell 2018)
Our Rts1 AID data

**Figure S1: Optimization and validation of AID phosphoproteomics method.** *A,* effects of bulk SCX pre-enrichment of phosphopeptide pool in reducing background of unmodified peptides. *B,* representative effect of SCX pre-enrichment on the final fraction of phosphopeptides after PolyMAC step. Identified phosphopeptides and unmodified peptides from LC-MS/MS were used to create the bar graph. *C,* representative distribution of phosphopeptides and unmodified peptides in the C18 high pH reverse phase fractions used for LC-MS/MS analysis. *D*, the indicated strains were grown to saturation in YPAD, serial diluted and spotted on YPAD agar plates supplemented with IAA and/or NaCl as indicated. Images were taken after 48 hours (-NaCl) or 96 hours (+ 1M NaCl) growth at 30 °C. *E*, IAA concentration-dependence of Rts1-ABD degradation was monitored by anti-V5 immunoblotting after 60-minute treatment of log-phase YPAD cultures. G6PDH is a loading control. "Rel. amt." is the load-normalized percent Rts1-ABD remaining relative to the mock treatment. *F*, degradation of Rts1-ABD in the large-scale nocodazole treatment phosphoproteomics experiment was confirmed by anti-V5 immunoblotting. *G,* fraction of large-budded cells after nocodazole arrest and either IAA or mock (DMSO) treatment was measured by microscopy. Data from a single culture are shown, which was representative of all biological replicates. *H,* comparison of overlap between our proteins with Rts1-dependent upregulated phosphosites and those identified from *rts1Δ* strains by Zapata et al. (53) and Touati et al. (78).

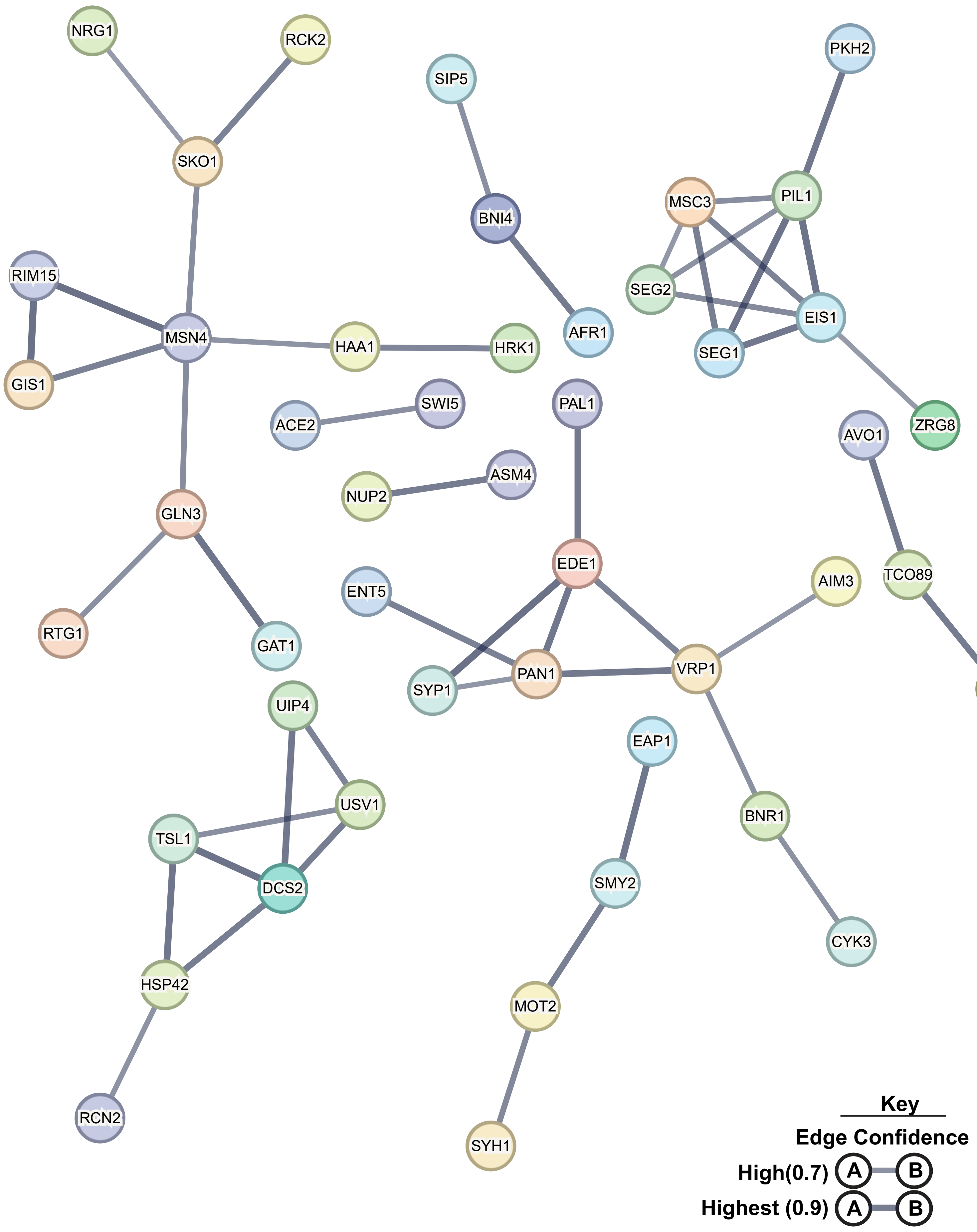
NRG1
RCK2
SIP5
PKH2
SKO1
BNI4
MSC3
PIL1
RIM15
SEG2
EIS1
MSN4
HAA1
HRK1
AFR1
SEG1
GIS1
SWI5
PAL1
ACE2
AVO1
ZRG8
ASM4
NUP2
GLN3
EDE1
ENT5
AIM3
TCO89
RTG1
GAT1
SYP1
PAN1
VRP1
PIB2
UIP4
EAP1
USV1
TSL1
BNR1
DCS2
SMY2
CYK3
HSP42
MOT2
RCN2
SYH1
Key
Edge Confidence
High(0.7)
A
B
Highest (0.9)
A
B

**Figure S2: STRING protein functional network.**

The STRING (54) protein functional interaction network predicted from the 111 upregulated phosphoproteins from the Rts1 AID phosphoproteomics dataset.

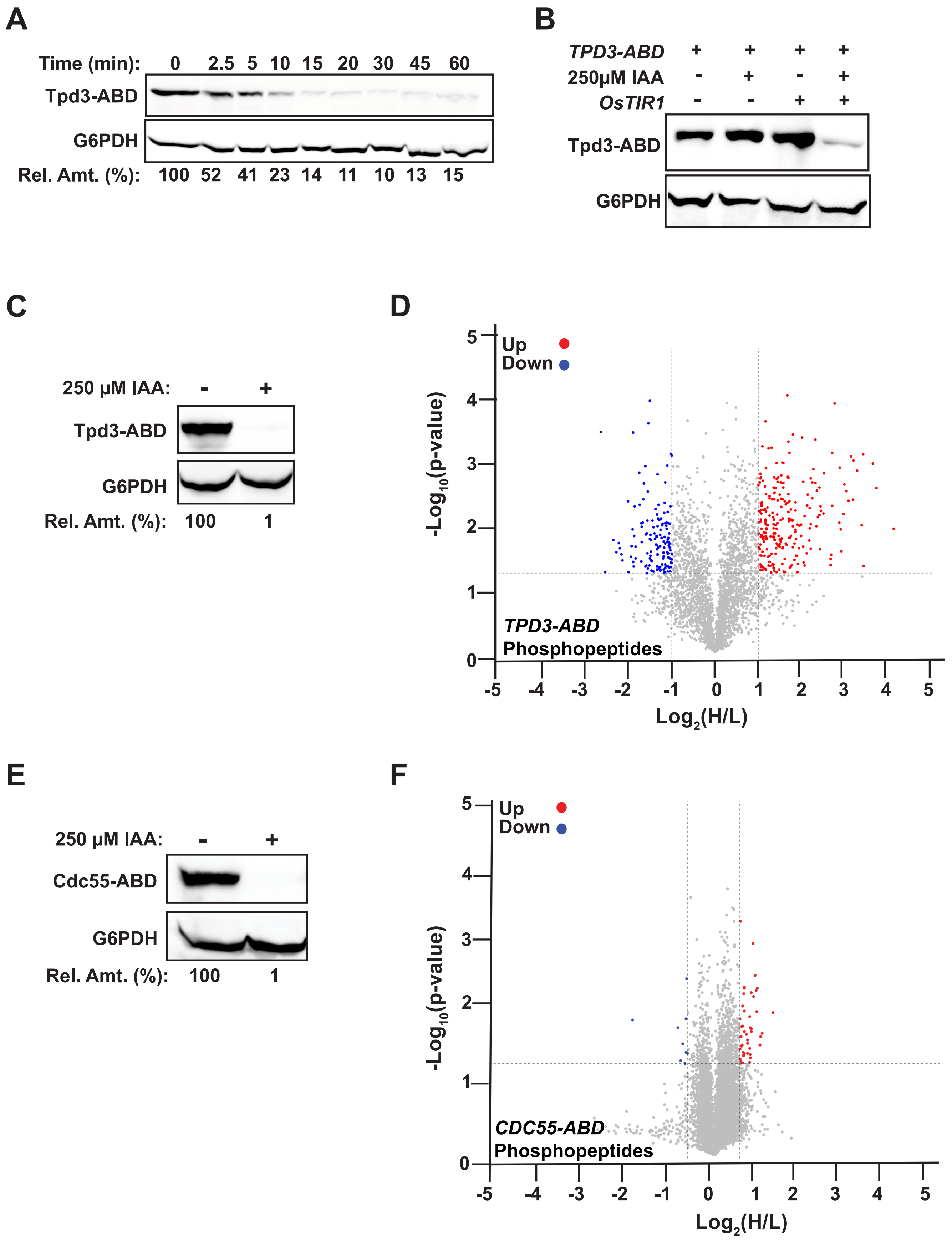
A
Time (min): 0 2.5 5 10 15 20 30 45 60
Tpd3-ABD
G6PDH
Rel. Amt. (%): 100 52 41 23 14 11 10 13 15
B
TPD3-ABD + + + +
250µM IAA - + - +
OsTIR1 - - + +
Tpd3-ABD
G6PDH
C
250 µM IAA: - +
Tpd3-ABD
G6PDH
Rel. Amt. (%): 100 1
D
Up
Down
-Log10(p-value)
Log2(H/L)
TPD3-ABD
Phosphopeptides
E
250 µM IAA: - +
Cdc55-ABD
G6PDH
Rel. Amt. (%): 100 1
F
Up
Down
-Log10(p-value)
Log2(H/L)
CDC55-ABD
Phosphopeptides

**Figure S3: Mitotic AID phosphoproteomic analysis of *TPD3-ABD* and *CDC55-ABD*.** *A,* Degradation kinetics of Tpd3-ABD in log phase YPAD cultures was monitored by anti-V5 immunoblotting after treatment with 250 μM IAA. G6PDH is a loading control. "Rel. amt." is the load-normalized percent Tpd3-ABD remaining relative to time 0. *B*, dependence of Tpd3-AID degradation on both IAA and *OsTIR1* was measured by immunoblotting as in A. *C,* confirmation of Tpd3-ABD degradation in the large-scale nocodazole phosphoproteomic experiment by anti-V5 immunoblotting. *D,* volcano plot for phosphopeptide H/L ratios from the *TPD3-ABD* nocodazole experiment. Cutoff threshold for Tpd3-dependent regulation was $-\text{Log}_{10}$(p-value) > 1.3 (p-value ≤ 0.05) and H/L cutoff +/- 1.0 (2-fold change). *E,* confirmation of Cdc55-ABD degradation in the large-scale nocodazole phosphoproteomic experiment by anti-V5 immunoblotting *F,* volcano plot for phosphopeptide H/L ratios from the *CDC55-ABD* nocodazole experiment. Cutoff threshold for Cdc55-dependent regulation was $-\text{Log}_{10}$(p-value) > 1.3 (p-value ≤ 0.05) and H/L cutoff +/- 0.585 (1.5-fold change). Volcano plot individual p-values were determined by t-test in Perseus. Red = upregulated peptides; blue = downregulated peptides; grey = non-regulated peptides.

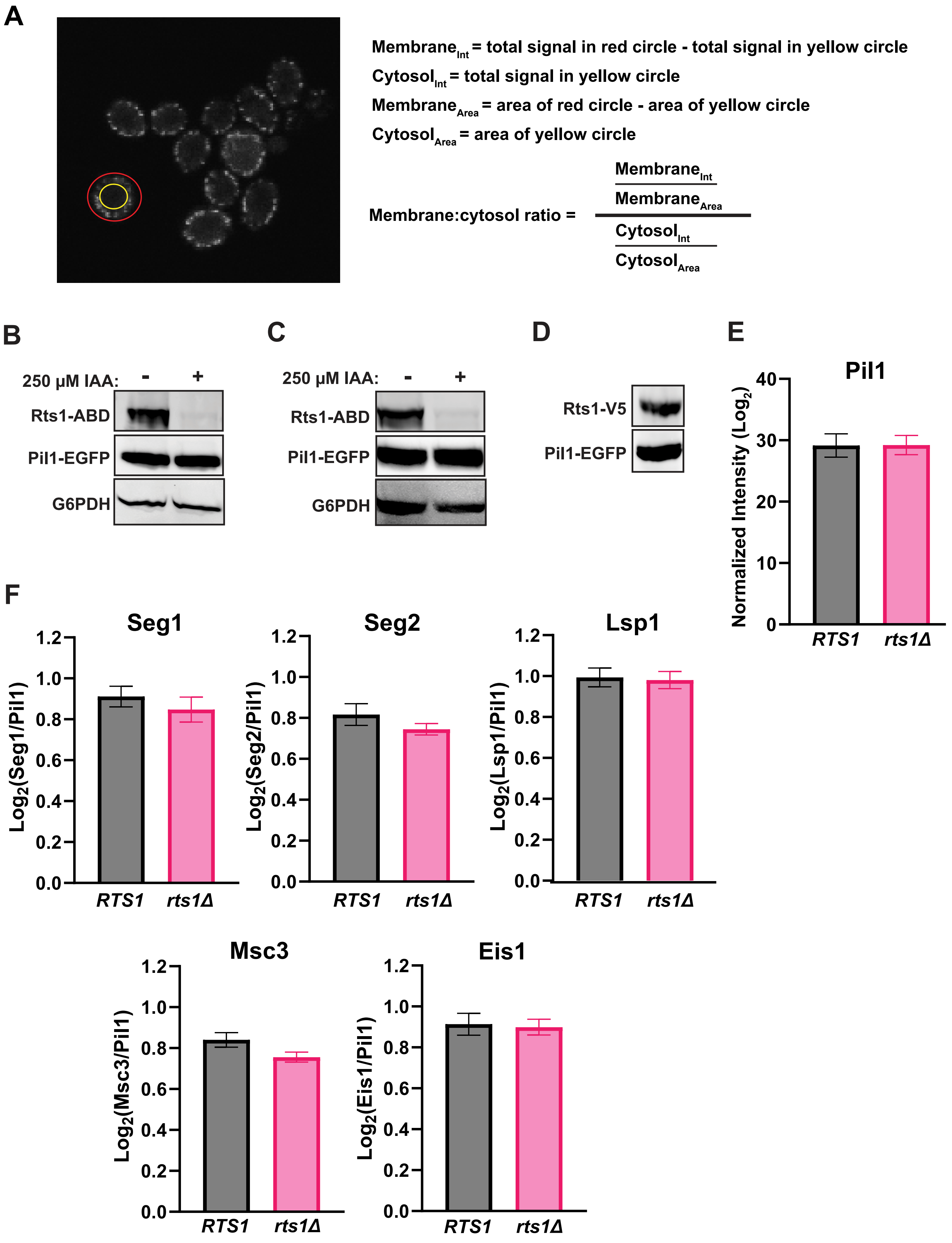
A
Membrane_Int = total signal in red circle - total signal in yellow circle
Cytosol_Int = total signal in yellow circle
Membrane_Area = area of red circle - area of yellow circle
Cytosol_Area = area of yellow circle
Membrane:cytosol ratio = (Membrane_Int / Membrane_Area) / (Cytosol_Int / Cytosol_Area)
B
250 µM IAA: - +
Rts1-ABD
Pil1-EGFP
G6PDH
C
250 µM IAA: - +
Rts1-ABD
Pil1-EGFP
G6PDH
D
Rts1-V5
Pil1-EGFP
E
Pil1
Normalized Intensity (Log2)
0
10
20
30
40
RTS1
rts1Δ
F
Seg1
Log2(Seg1/Pil1)
Seg2
Log2(Seg2/Pil1)
Lsp1
Log2(Lsp1/Pil1)
Msc3
Log2(Msc3/Pil1)
Eis1
Log2(Eis1/Pil1)
0.0
0.2
0.4
0.6
0.8
1.0
1.2
RTS1
rts1Δ

**Figure S4: Supporting information for PP2A$^{Rts1}$ regulation of Pil1 localization and eisosome subunit interactions.**

*A,* method used to quantify membrane:cytosol Pil1-EGFP fluorescence ratio for all fluorescence microscopy experiments. The intensity (Int) per unit area was determined for the membrane region and the cytosol region in each cell body using ImageJ software and the indicated equations. The ratio of these two values is independent of cell-to-cell and image-to-image variation and therefore serves as a useful value for comparing membrane-associated Pil1-EGFP signal between different strains and conditions. *B-C,* confirmation of Rts1-ABD degradation for Pil1-EGFP localization experiment in metaphase arrest (B), and asynchronous log phase culture (C) experiments. Rts1-ABD was monitored by anti-V5 immunoblotting, Pil1-EGFP by anti-GFP immunoblotting, and G6PDH is a loading control. *D*, expression confirmation of *RTS1-3xV5* from centromeric complementation plasmid in *rts1Δ PIL1-EGFP* by anti-V5 immunoblotting. *E,* Comparison of summed Pil1 peptide intensities from Pil1-EGFP IP from *RTS1 and rts1Δ* strains. *F,* Comparison of Pil1-associated eisosome proteins from Pil1-EGFP IP-MS analysis. The summed intensities of peptides from each detected eisosome-associated protein were divided by the summed Pil1-EGFP peptide intensity and plotted. A t-test was used to compare *RTS1* vs. *Δrts1, and the differences were not statistically significant ($p \geq 0.05$) in each case*.

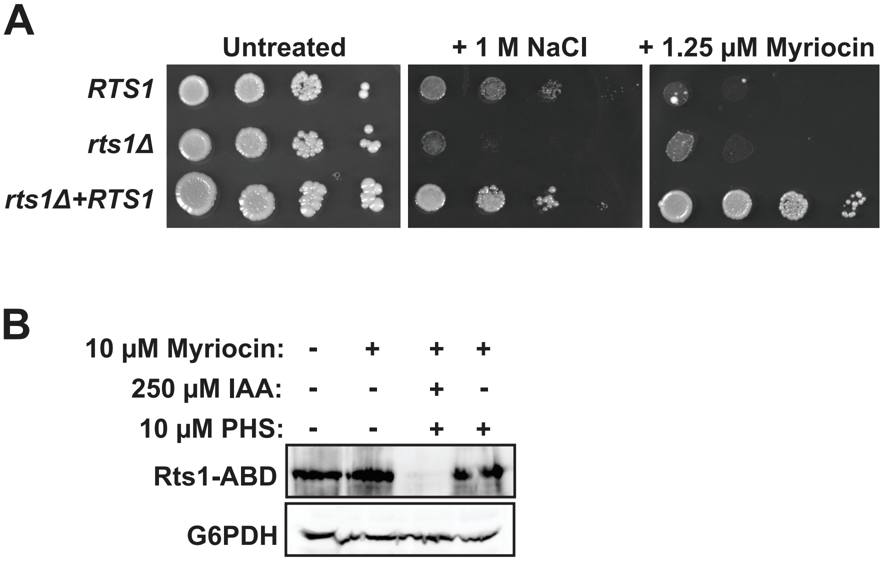


**Figure S5: Supporting information for PP2ARts1 regulation of Pil1 localization in sphingolipid metabolism.** A, Serial dilution spot assay in which the indicated strains were grown to saturation in YPAD, serially diluted, and spotted onto YPAD agar plates with the indicated supplements. Growth was monitored at 30°C for 96 hours. B, Degradation confirmation anti-V5 immunoblot from the microscopy experiment shown in Fig. 4B.
